# Melanocortin 4 receptor-expressing neurons in the lateral stripe of the striatum are involved in affect regulation and motor control

**DOI:** 10.1101/2024.05.31.596779

**Authors:** Johan Sköld, Gisela Paola Lazzarino, Myra Nett, Joost Wiskerke, David Engblom

**Affiliations:** Center for Social and Affective Neuroscience, Department of Biomedical and Clinical Sciences, Linköping University, Linköping, Sweden

**Author notes:** These authors contributed equally and share first authorship (Johan Sköld, Gisela Paola Lazzarino).

**Keywords:** Aversion, Reward, Salience, Motor activity, in vivo calcium imaging, Optical intracranial self-stimulation, Fear conditioning, rsChrmine, dLight, iGluSnFR, Open field test, Real-time place preference test, Wheel running

## Abstract

The dopaminergic system is crucial for affect regulation. Melanocortin 4 receptors (MC4R) in the ventral striatum have been shown to be necessary for establishing aversive states. Here, we set out to functionally characterize MC4R-expressing striatal neurons. MC4Rs were enriched in atypical Dopamine receptor 1 (D1) neurons in the lateral stripe of the striatum (LSS), an understudied area in the ventrolateral striatum. Fiber photometry recordings showed that MC4R neuron activity and local dopamine release in the LSS were increased by rewarding as well as aversive stimuli. Moreover, MC4R neuronal activity and glutamate release correlated strongly to body movement. Optogenetic activation of MC4R-LSS neurons was rewarding in a real-time place preference test and a self-stimulation paradigm, increased locomotor activity and induced striatal dopamine release. Collectively, our findings suggest that MC4R-LSS neurons are activated by salient stimuli of both rewarding and aversive character and that they induce positive affect, dopamine release and locomotion.

## Introduction

Understanding the neurobiological mechanisms driving negative affect is crucial for developing new treatment strategies for a wide variety of pathological conditions, from major depressive disorder to negative affect in inflammatory diseases. The dopaminergic system, including the striatum, is involved in the induction of negative affect (Belujon & Grace, 2017; Bromberg-Martin et al., 2010; de Jong et al., 2019; Klawonn & Malenka, 2018; Tye et al., 2013). The melanocortin system has been identified as a key regulator of motivated behaviors within the ventral striatum (Pandit et al., 2015). It has been shown that signaling through the melanocortin 4-receptor (MC4R) in dopamine receptor 1 (D1) neurons of the nucleus accumbens (NAc) is increased in anhedonia (Lim et al., 2012). Moreover, our laboratory has previously demonstrated that MC4Rs in the striatum are crucial for establishing aversive responses. We found that mice lacking MC4Rs were seeking, instead of avoiding, environments associated with aversive stimuli, indicating that they found the aversive stimuli rewarding. When MC4R expression was rescued in the striatum, or in D1 expressing neurons, of MC4R -/- mice, the mice displayed normal responses to aversive stimuli. This suggests that MC4Rs in striatal D1 neurons work as a switch controlling the valence of aversive stimuli (Klawonn et al., 2018), but the mechanisms behind the “switching” are unclear and the identity of the MC4R expressing neurons in the striatum has not been elucidated in any detail. A brain-wide mapping study in mice suggests that MC4R expressing neurons are enriched in the lateral stripe of the striatum (LSS; Liu et al., 2003), an often-overlooked structure located in the ventrolateral striatum. Interestingly, recent single-cell RNA-sequencing studies indicate that the LSS is also enriched in a population of neurons that does not cluster with the classic D1- or D2-neurons. These cells are defined alternately as “hybrid D1 neurons”, “eccentric spiny projection neurons” or “atypical D1 neurons” (Chen et al., 2021; Gokce et al., 2016; Saunders et al., 2018; Stanley et al., 2020).

Here, we characterized the identity and functions of the MC4R-expressing neurons located in the LSS, showing that these neurons share characteristics with atypical spiny projection neurons. Using transgenic mice and optical techniques, such as in vivo calcium imaging and optogenetics, combined with behavioral analysis, we show that the MC4R-LSS neurons are activated by both rewarding and aversive stimuli, and their activation is rewarding.

## Materials and methods

### Animals

Experimental protocols were conducted according to international and national guidelines for animal research and were approved by the Research Animal Care and Use Committee in Linköping, Sweden.

All experiments were done using mice aged between 8-20 weeks at the onset of experiments. Animals were kept in a pathogen-free facility on a regular 12-hour light/dark cycle (lights off 07.00 p.m.) and a temperature of 22 ± 2°C in individually ventilated cages. All experiments were performed in the light phase. Food and water were provided *ad libitum*, except otherwise stated. Mc4rtm3.1(cre)Lowl/J (Mc4r-cre, #030759), B6.Cg-Gt(ROSA)26Sortm9(CAG-tdTomato)Hze/J (Ai9, #007909), B6.Cg-Tg(Mc4r-MAPT/Sapphire)21Rck/J (Mc4r-GFP, #008323), B6.Cg-Tg(Drd1a-tdTomato)6Calak/J (Drd1-tdTomato, #016204), 129S6.FVB(B6)-Tg(Drd1a-cre)Agsc/KndlJ (D1-cre, #028298) and Tg(Pomc1-cre)16Lowl/J (POMC-cre, #005965) lines have previously been described in the literature (Ade et al., 2011; Balthasar et al., 2004; Garfield et al., 2015; Liu et al., 2003; Madisen et al., 2010), and were purchased from The Jackson Laboratory. MC4R-tdTomato animals were obtained by crossing Mc4-cre and Ai9 lines, and Drd1-tdTomato-MC4R-eGFP line was obtained by crossing Mc4r-GFP and Drd1-tdTomato. All mice used in this study had a C57BL/6 background, except for POMC-Cre (mixed FVB/N and C57BL/6J strains). Animals were assigned into different experimental groups balanced by sex and age, and single housed after surgeries. We did not observe any noticeable differences in the results between male and female mice. As a result, data obtained from both sexes were combined.

### Viral constructs

The following viruses used in the current study were obtained from AddGene: pAAV5-EF1a-doublefloxed-hChR2(H134R)-EYFP-WPRE-HGHpA (20298-AAV5; hChR2), pAAV5-Ef1a-DIO-EYFP (27056-AAV5; EYFP), pAAV5-EF1a-double floxed-hChR2(H134R)-mCherry-WPRE-HGHpA (20297-AAV5; hChR2-mCherry), pAAV1-Ef1a-fDIO mCherry (114471-AAV1; mCherry), pAAV5-flex-taCasp3-TEVp (45580-AAV5; Casp3), pGP-AAV9-syn-FLEX-jGCaMP8m-WPRE (162378-AAV9; GcaMP8m), pAAV5-hSyn-dlight1.2 (111068-AAV5; dlight1.2) AAV9.CamKII.GcaMP6s.WPRE.SV40 (107790-AAV9; GcaMP6s). ssAAV-1-hSyn1-chI-dlox-SF_iGluSnFR(A184S)(rev)-dlox-WPRE-SV40p(A) (v357-1; SF_iGluSnFR) was obtained from the Zürich Viral Vector Facility. pAAV8-Ef1a-DIO-rsChRmine-oScarlet-WPRE (rsChRmine) and pAAV8-Ef1a-DIO-mScarlet-WPRE (mScarlet) were obtained from the Deisseroth Lab via the Stanford Gene Vector and Virus Core (RRID:SCR_023250).. All vector titers ranged from 8.4×10^12^ to 2.3×10^13^ viral molecules/ml.

### Stereotactic injections and optic fiber implantation

For all stereotactic surgeries, mice were first administered a subcutaneous injection of buprenorphine (0.01 mg/kg; Vetergesic vet; Orion Pharma) as an analgesic 30 min prior to intervention. Subsequently, mice were anesthetized with 3% isoflurane, placed in a stereotaxic apparatus (David Kopf Instruments), and maintained at a constant isoflurane level of 1.0–1.5% throughout the surgical process. Adeno-associated virus (AAV) vectors were unilaterally injected at a rate of 100 nL/min using a gastight Hamilton Neuros syringe (33G), an UMP3 micro-syringe injector and a Micro4 controller (World Precision Instruments Inc.). All injections (300 nL) were performed in the LSS using the following coordinates from Bregma: Anterior/posterior [AP]: 1.1; medial/lateral [ML]: ±2.0; dorsal/ventral [DV]: −4.7 for fiber photometry experiments, and [DV]: −4.9 for optogenetic experiments. For projection mapping analysis, 100 nL of pAAV5-Ef1a-DIO EYFP was injected in the LSS of MC4r-cre mice using the aforementioned coordinates, or in the arcuate nucleus (ARC) of POMC-cre animals, using the coordinates AP: −1,5; ML: ±0,5; DV: −5,8. To induce neuronal ablation, 300 nL of a mixture of pAAV5-flex-taCasp3-TEVp with pAAV5-Ef1a-DIO-EYFP (9:1) was bilaterally injected into the LSS of MC4r-cre mice or their wild type littermates (control group). For experiments involving optogenetic activation combined with calcium imaging, MC4r-cre animals were injected in the LSS, employing the same coordinates as stated above, with a 1:1 mixture of pAAV5-hSyn-dlight1.2 and pAAV8-Ef1a-DIO-rsChRmine-oScarlet-WPRE (active opsin) or pAAV8-Ef1a-DIO-mScarlet-WPRE (control group). The following coordinates from Bregma were used for optogenetic activation of NAcC (AP: 1,1; ML: ±1,5; DV: −4,1), medial nucleus accumbens shell (AP: 0,8; ML: ±0,5; DV: −4,1), and dorsal striatum (AP: 0,8; ML: ±1,6; DV: - 3,0). For fiber photometry in the NAcC the following coordinates were used: AP: 1,1; ML: ±1,5; DV: −4.1. For all virus injections, the injection needle was left in place for 10 min after injection to ensure proper diffusion. Following viral injections, animals were either implanted with an optic fiber implant or surgically closed with sutures. For optogenetic experiments, optic fiber implants (Doric Lenses Inc.; O.D. 5 mm, Fiber Core 200/250 μm, NA 0.66) were positioned 300 μm above the injection sites. For fiber photometry, optic fibers (Doric Lenses Inc.; O.D. 5 mm, Fiber Core 400/430 μm, NA 0.66) were placed 100 μm above the targeted region. Optic fibers were fixed to the skull using dental adhesive resin cement (Super-bond C&B, Sun Medical co.) and black acrylic resin (Ortho-Jet, Lang Dental Manufacturing Co.). Analgesic treatment with Carprofen (5 mg/kg; Rimadyl, Pfizer) was administered during the first 48 hours post-surgery. Animals were allowed to recover for at least 3 weeks after surgery to allow viral expression before conducting experiments. Prior to fiber-photometry experiments, the signal was screened and mice with very low signal-to-noise ratio were excluded. Following the behavioral experiments, construct expression, injection site, and fiber location were confirmed through fluorescent immunohistochemistry.

### Fiber photometry

All fiber photometry measurements were conducted using hardware from Tucker-Davies Technologies (TDT). Excitation of the fluorophores (GcaMP8m, GcaMP6s or dLight1.2) was achieved using 465 nm (approximately 50 ± 10 µW) and 405 nm (approximately 25 ± 5 µW) light, with different modulation frequencies. Raw data were demodulated in real-time using Synapse (TDT) into the active, 465 nm-excited channel, and the isosbestic 405 nm-excited control channel. TTL pulses from Med Associate boxes were recorded for aligning the recordings to specific events. Data analysis was performed using fpExplorer (unpublished), an in-house developed fiber photometry analysis software. The change in fluorescence relative to an initial baseline fluorescence (dF/F) was calculated using the Barker method (Bruno et al., 2021).

### Behavioral paradigms

Mice were habituated to handling and connection to a fiber patch cord for several days prior to the different test sessions.

### Pavlovian fear-conditioning

For fear conditioning, mice were tethered to the photometry patch cable, and placed in a mouse conditioning chamber (Med Associates Inc.). Animals were exposed to a 30s CS tone, and during the last 2s a 0.5 mA foot shock was delivered via the grid floor of the conditioning chamber. Animals were exposed to 6 trials. Inter-trial interval time was quasi-random with a mean of 3 min. On the following day, fear memory was tested by exposing the animals to 6 × 30s cue tones without foot shocks, in the same conditioning chamber. Mice were videotaped during the conditioning and test session allowing quantification of body movement using the “Activity analysis” feature of EthoVision XT (Noldus). The percentage of pixels changed per frame was measured. Calcium activity or dopamine release was measured continuously throughout the entire session. Activity was correlated to the fluorescence signal with the MatLab function “xcorr”.

To compare the neural response with and without D1-receptor-antagonism, animals were pretreated with either saline or SCH23390 hydrochloride (0.2 mg/kg, Tocris, Cat. No. 0925) dissolved in physiological saline and injected i.p. in a volume of 4 µL/g 20 min prior to fear conditioning described above. The following day, the animals were retested with the same conditions after pretreatment with the compound not administered on day 1. Maximal calcium transient height during the 2s shock was measured (in dF/F) and compared between the 2 treatments.

### Combined reward and aversion

Animals with GcaMP8m in MC4R-LSS neurons or dLight in the LSS area were trained in a Pavlovian conditioning where a 5s tone predicted a sugar pellet delivery (20 mg; Bio-Serv). Med Associates boxes were used together with custom, 3D-printed food-ports for allowing video recording of delivery and consumption of sugar pellets while the mice were attached to the fiber optic cable. The mice were food restricted to approximately 50% of their daily food intake for 3 nights before training started and kept at 85-90% of their initial body weight throughout the experiment. Mice were exposed to 5 training sessions consisting of 30 trials with quasi-randomized inter-trial intervals with a mean of 60s. On the sixth day, 10 of the 30 sugar pellet deliveries were unpredicted. On day 7, 10 of the trials were predicted by the tone but the pellet delivery was omitted. On day 8, the session started with 10 predicted pellet deliveries. The house light was then turned off and 10 foot-shock trials predicted by 5s onset of cue-lights started. To compare the magnitude of GcaMP/dLight responses in response to reward compared to aversion, maximal dF/F within 10s from pellet delivery/foot-shock was measured. The mean peak dF/F was calculated for each condition for each animal. Paired comparisons of responses to different conditions were then performed.

### Intracranial self-stimulation

Mice were connected to an optical fiber linked to a 460 nm LED source (Prizmatix Ltd.) and tested in a standard mouse operant chamber (Med Associates Inc.). The chamber light was turned on to indicate the start of the test. The operant chamber contained two nose-poke holes that were illuminated during the whole test. One of these holes was randomly paired to a 2s blue light stimulation (20 Hz, 10ms pulses, approximately 12 mW into the brain). When mice poked into the active hole, the port illumination ceased while the animal received the stimulation. Responses on the active hole during the 2 s stimulation period were recorded but did not lead to any additional stimulation. If the animal poked in the inactive hole, no stimulation was delivered, and the port remained illuminated. Each session lasted for 30 min and was performed for 3 consecutive days. The designation of the active port was changed daily.

### Real-time place preference

Real-time place preference test was conducted one week after the completion of the intracranial self-stimulation test. A 3-chambered Panlab Spatial Place Preference Box (Harvard Apparatus) was used. On day 1, a pretest was conducted. Mice were connected to an optical fiber and allowed to move freely between the chambers of the box, to establish any specific preference. The chamber in which each mouse spent a lesser amount of time was designated as the non-preferred chamber. This specific non-preferred chamber was then coupled with blue light stimulation (20 Hz, 10ms pulses, approximately 12 mW into the brain) during the subsequent days of testing (days 2-4). For the testing phase (days 2-4), animals were attached to an optical fiber connected to a LED source. The animals were placed within the corridor of the apparatus and permitted to move freely amongst the chambers of the box. The animals received stimulation when in the light-paired side, and all sessions were conducted for 15 min. EthoVision XT tracking software (Noldus) controlled animal location, and light stimulation in real-time Time spent in each chamber was measured to elucidate if the optogenetic activation establishes place-preference or aversion.

### Locomotor activity

After the completion of the real-time place preference test, mice were tested for locomotor activity. For that, the animals were attached to an optical fiber connected to a LED source and placed in the center of an open-field arena (45 cm × 45 cm × 40 cm). Locomotion was assessed for 15 min: an initial 5 min free exploration without stimulation, followed by 5 min of 20 Hz blue light stimulation (10ms pulses, approximately 12 mW into the brain), concluded by 5 min of no stimulation.

Spontaneous locomotor activity was also evaluated in MC4r-cre mice with neuronal ablation (n=8 ablated, 8 controls), pre- and post-injection of casp3. Animals were placed in an open-field arena (45 cm × 45 cm × 40 cm) for 10 min.

Animals were monitored using a camera and EthoVision XT tracking software (Noldus). Total distance traveled was used to evaluate the effect of neural activation or ablation.

### Voluntary wheel-running

Voluntary wheel-running activity was evaluated over two distinct time frames: a 10-day period prior to Casp3 injection and a subsequent 10-day period 3 weeks post-injection. Animals (n=8 ablated, 8 controls) were single housed in filtertop cages (37 x 21 x 18 cm) and given continuous access to wireless running wheels. The quantification of wheel running behavior was conducted by tracking the number of wheel rotations over time. Data was exported and expressed as cumulative turns during each test period. During the whole experiment, food intake and body weight were measured at 3–4-day intervals. Measured food intake was divided by measurement interval and expressed as food intake per day.

### Combined optogenetic and fiber photometry

Four weeks after surgeries, animals were placed in an operant chamber and connected to the optical fiber. During fiber photometry recordings, red light (625 nm, 20 Hz, 10ms pulse width, 5 mW from fiber tip) was delivered through the same fiber cable at either 2, 5, 10, 20 or 40 pulses (randomly chosen) with a total of 10 trials per condition, 50 trials in total. To quantify the dopamine released in response to the optical stimulation, area under the curve of the dF/F was measured and compared between the conditions and the active/control group. The following week, light intensity was titrated. This time, animals received 4 blocks of 10 stimulation trials (45s intertrial interval). In each trial, 5 pulses of red light (625 nm, 20 Hz, 10ms pulse width) were delivered, with the light intensity varying between trial blocks: 1 mW, 5 mW, 10 mW and 15 mW, measured from the fiber tip. The order of light intensities was randomly assigned to each animal. For each trial, the peak dF/F value following stimulation onset was identified. Next, per animal, the median of these peak values was derived for each of the 4 trial blocks. These medians were then used to compare the different light intensities.

### Ex-vivo slice opto-photometry

Animals were bilaterally injected with a mixture of GcaMP8m or dLight1.2 combined with rsChrmine or mScarlet (rsChrmine in one hemisphere, mScarlet in the other hemisphere). At least 3 weeks later, animals were then anesthetized with 4% isoflurane. The brain was removed and transferred to ice-cold cutting solution bubbled with 95% O_2_ and 5% CO_2_. Content of cutting solution (in mM): 92 NMDG, 20 HEPES, 25 glucose, 30 NaHCO_3_, 1.2 NaH_2_PO_4_, 2.5 KCl, 5 sodium ascorbate, 3 sodium pyruvate, 2 thiourea, 10 MgSO_4_, and 0.5 CaCl_2_, pH 7.4. Coronal sections (300 μm) were cut using a vibratome (Leica). Sections were transferred to 35°C aCSF solution for recovery (in mM): 125 NaCl, 2.5 KCl, 1.25 NaH_2_PO_4_, 1 MgCl_2_, 11 glucose, 26 NaHCO_3_, 2.4 CaCl_2_ (310 mOsm, pH 7.4) bubbled with 95% O_2_, 5% CO_2_. The aCSF with sections were placed at room temperature at least 30 min prior to recording. Sections were put in a recording chamber and perfused with fresh aCSF at a rate of 2 mL/min. Using a fluorescent microscope, transfected cells could be visualized. An optic fiber was placed above the fluorescent cells and fiber photometry recordings were performed as previously described. To stimulate rsChrmine, 5 pulses of red light (625 nm, 20 Hz, 10ms, 23 mW from fiber tip) was used. GcaMP or dLight responses were measured during optical stimulation. Data was processed as described above.

### Immunofluorescence

Animals were euthanized in CO_2_, transcardially perfused with 0.9% saline solution followed by 4% paraformaldehyde (PFA) in PBS (pH 7.4). Brains were removed, postfixed overnight in a 4% PFA solution, then transferred to a 30% sucrose-PBS solution until sinking. Brains were frozen in −80°C isopentane. Coronal sections of 40 μm were cut using a cryostat (Leica CM1959, Leica), then placed into a cryoprotectant buffer (composed of 0.1 M phosphate buffer, 30% ethylene glycol, and 20% glycerol), and stored at −20°C until further use. For free-floating immunofluorescent labeling, the sections were washed in PBS, followed by an incubation with blocking solution (1% BSA and 0.3% Triton X-100 in PBS). Subsequently, the sections were incubated overnight with primary antibodies diluted in blocking solution: rabbit anti-foxP2 (1:2000; Abcam, ab16046), rabbit anti-red fluorescent protein (anti-RFP, 1:1000; MBL International, PM005), chicken anti-GFP (1:10000; Abcam, ab13970). The sections were then washed in PBS and incubated for 2h with appropriate secondary fluorescent antibodies Alexa Fluor 488/568 (all provided by Invitrogen): anti-rabbit (1:500; A21206; or 1:1000; A10042), or anti-chicken (1:1000; A11039). Finally, the sections were washed again with PBS, and mounted on glass slides using ProLong Glass Antifade Mounting agent (Thermofisher). Cells expressing MC4r-tdTomato were visualized without the use of immunohistochemistry. Sections were analyzed using a confocal microscope (Zeiss LSM 700, Carl Zeiss AG) or a widefield microscope (Leica Dmi8) with 405, 488, 555, and 639 nm diode lasers. Confocal pictures were used for manual counting of co-expression of FOXP2 and D1 in MC4R-LSS neurons.

### RNA-Scope fluorescent in situ hybridization

Animals were transcardially perfused, and their brains removed, post-fixed, and cryoprotected in 30% sucrose, as described above. 16 µm brain sections were cut at the striatal level using a freezing microtome, mounted on SuperFrost Plus Gold slides (ThermoFisher), and baked at 60°C overnight. In situ hybridization was performed using RNAscope Multiplex Fluorescent assay v2 (Advanced Cell Diagnostics) following manufacturers’ protocol. The mc4r probe (Cat No. 319181) was purchased from Advanced Cell Diagnostics (Newark). Pretreatment target retrieval was done at 98–99°C for 7 min, and sections were treated with Protease Plus for 22 min. Opal 570 tyramide (1:2000; Akoya Biosciences) was used to detect the MC4R probe. Microphotographs were obtained using a widefield microscope (Leica Dmi8) at 25x magnification.

### Statistics

GraphPad Prism 10 was used to make graphs and perform statistical analyses.. Shapiro-Wilk tests were performed to evaluate the normal distribution of the data. Two tailed paired t-tests and one sample t-tests were used for data with equal variances in the groups. For repeated measures, repeated measures ANOVA followed by Šídák’s multiple comparisons test or multiple Mann-Whitney tests with false discovery rate were performed, as appropriate. Statistical tests used, as well as sample sizes, are stated in figure legends. Some data is descriptive and does not require statistical comparations. Data is presented as mean, and error bars show SEM.

## Results

### MC4R is expressed by atypical medium spiny neurons in the lateral stripe of the striatum

To visualize the localization of MC4R-expression in the striatum, we crossed MC4R-cre mice with mice carrying a floxed tdTomato gene. We found that the MC4R-expression is enriched in the LSS, while expression in the rest of the striatum is sparse (figure 1A). Expression of MC4R in adult mice was confirmed using in situ hybridization (supplementary figure S1A). In a mouse expressing GFP under the MC4R-promoter and tdTomato under the D1-promoter, we found that the majority of MC4R-LSS neurons also expressed D1-receptors (figure 1B). Since the LSS is known to harbor atypical D1-neurons with high FOXP2-expression (Chen et al., 2021; Gokce et al., 2016; Saunders et al., 2018; Stanley et al., 2020), we counterstained sections from MC4R-tdTomato brains for FOXP2. We found high FOXP2-expression in the LSS, and co-expression of FOXP2 in MC4R-LSS neurons (figure 1C), with 96,4% (1207/1252) of MC4R-LSS neurons expressing FOXP2. To investigate projection patterns of MC4R-LSS neurons, we injected a cre-dependent EYFP vector into the LSS of MC4R-cre mice. Projection mapping of these mice showed dense projections to the substantia nigra pars reticulata, among other brain areas (figure 1D-E and supplementary figure S1B-F). To investigate if the MC4R-LSS neurons receive afferents from POMC-neurons in the arcuate nucleus of the hypothalamus (ARC; the primary source of the MC4R-agonsist α-melanocyte stimulating hormone, α-MSH), we injected a cre-dependent EYFP vector in the ARC of POMC-cre mice. Labeled fibers were seen in the medial nucleus accumbens shell, but not in the LSS, indicating that MC4R-LSS neurons do not receive direct synaptic α-MSH inputs from the ARC (supplementary figure S2). Altogether, these results indicate that the MC4R is expressed by atypical D1-neurons in the LSS, and that the MC4R-cre line can be used to specifically study atypical D1-neurons.

**Figure 1.**
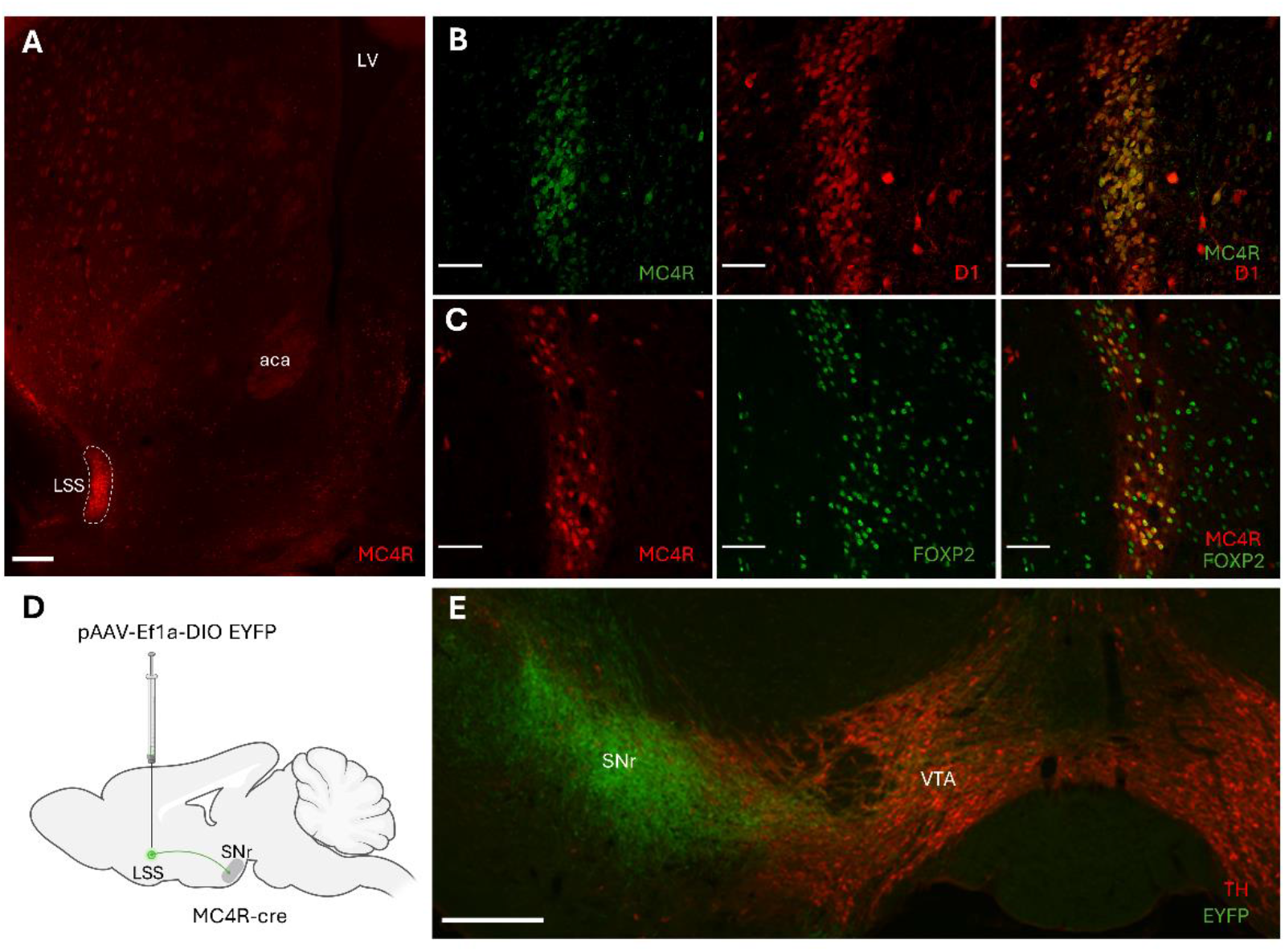
MC4R is expressed by substantia nigra-projecting atypical D1-neurons in the lateral stripe of the striatum. (A) Coronal section of an MC4R-cre, tdTomato-mouse showing the expression pattern of MC4R in the striatum. MC4R-expressing neurons are enriched in the lateral stripe of the striatum (LSS) and sparse in the rest of the striatum. (B) Confocal picture showing colocalization (right) of MC4R-(green) and D1-expression (red) in the LSS of a mouse expressing GFP under the MC4R-promoter and tdTomato under the D1-promoter. (C) MC4R-LSS neurons co-express FOXP2 (right), a marker for atypical D1-neurons. MC4R-cre-tdTomato (red) and FOXP2 (green) are expressed by the same neurons. (D) Schematic depicting the site of the virus injection and the major projection from LSS to substantia nigra pars reticulata (SNr). MC4R-cre mice were injected with pAAV-Ef1a-DIO EYFP in LSS. (E) Representative image of the projections of LSS-MC4R neurons to SNr. Fibers from MC4R-LSS neurons are shown in green, and tyrosine hydroxylase (TH)-expressing cells in red. Scale bars 200 µm (A, E) and 50 µm (B, C). LV: lateral ventricle, aca: anterior commissure, VTA: ventral tegmental area.

### MC4R-LSS neurons are activated by aversive foot-shocks

Since the MC4R has been reported to be necessary for aversive signaling, we measured how the MC4R-LSS neurons respond to aversive stimuli. To test this, we injected a viral vector carrying a cre-dependent genetically encoded calcium indicator (GcaMP8m) into the LSS of MC4R-cre mice and implanted an optic fiber over the LSS, allowing us to measure intracellular calcium as a proxy for neural activity with fiber photometry. We found an increase in firing of MC4R-LSS neurons during exposure to an aversive foot-shock (figure 2A-D). In contrast, no apparent change in intracellular calcium was observed in response to the conditioned tone predicting the shock during the acquisition session or the subsequent day, when the shock was omitted (figure 2C).

**Figure 2.**
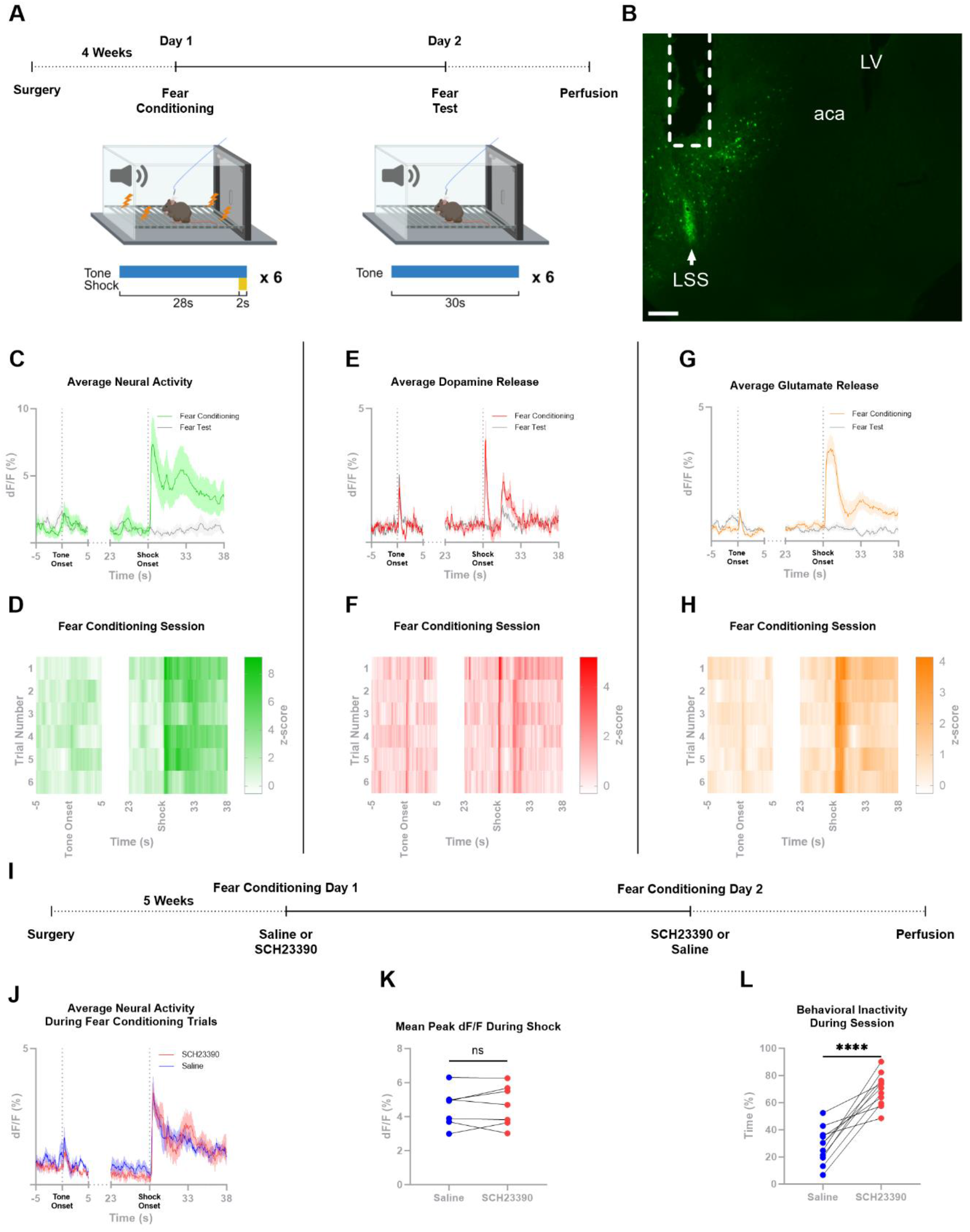
MC4R-LSS neurons are activated by aversive foot-shocks in a D1-independent manner. (A) Timeline of experiments. (B) Representative picture of GCaMP8m-expression and fiber placement in the lateral stripe of the striatum (LSS) of an MC4R-cre mouse. Fiber tract marked with dashed line; LSS marked with arrow. (C) GCaMP8m signal measured with fiber photometry from MC4R-LSS neurons shows activation of MC4R-LSS neurons in response to aversive shocks, but not to the onset of the predictive tone cue. No change in neural activity was seen in response to shock omission in the fear test on day 2. Mean and SEM from 6 mice, 6 trials per mouse. (D) Heatmap of the data shown in (C) but split out per trial showing that the neural response is similar across all trials. Mean from 6 mice. (E) Dopamine measurements from the LSS during the same behavioral paradigm as in (C). Dopamine is released in the LSS in response to both the aversive foot-shocks and their predictive tone cues. No clear change in dopamine release was seen in response to shock omission in the fear test on day 2. Mean and SEM from 7 mice, 6 trials per mouse. (F) Heatmap of the data shown in (E) but split out per trial showing that the tone-induced dopamine release develops during the session. Mean from 7 mice (G) iGluSNFR-measurements of glutamate release onto D1-expressing neurons in the LSS during fear conditioning. Glutamate is released onto D1-expressing neurons in response to the aversive foot-shock. A small release can be seen in response to the tone onset. Mean and SEM from 10 mice, 6 trials per mouse. (H) Heatmap of the data shown in (G), but split out per trial, showing that the shock-induced glutamate release is similar across trials and that the tone-induced release is developed within the session. Mean from 10 mice. (I) Timeline of the crossover experiment testing if shock-induced activity of MC4R-LSS neurons is D1-dependent. (J) Group mean peri-event traces from the experiment shows a robust increase in neural activity in response to the foot-shock after both saline-injection and pretreatment with the D1-antagonist SCH23390 (0.2 mg/kg; n=7). (K) Mean peak size during the shock for each animal showing that SCH23390 does not affect the shock-induced neural response (n=7, paired two-tailed t-test). (L) Analysis of the inactivity time during the sessions showing that mice spend significantly more time inactive after SCH23390 injection compared to saline injection (n=7, paired two-tailed t-test). *** p < 0.001. aca: anterior commissure; LV: left ventricle.

### Neurotransmitter release in the LSS during aversive fear conditioning

Two of the primary afferent neurotransmitters to striatal medium spiny neurons are dopamine from midbrain dopaminergic neurons and glutamate from the cortex and the thalamus(De Jong et al., 2022; Hunnicutt et al., 2016). In this study, we aimed to elucidate the afferent signals affecting MC4R-LSS neurons, by measuring the dopamine and glutamate release in the LSS during a fear conditioning paradigm, using fiber photometry, the dopamine sensor dLight1.2 and the glutamate sensor SF_iGluSnFR. To increase signal-to-noise ratio for the cre-dependent glutamate sensor, D1-cre instead of MC4R-cre animals were used. We found a robust and reproducible increase in dopamine release in response to both foot shocks and the conditioned tone predicting foot shocks (figure 2E-F), while glutamate was released primarily in response to the shock (figure 2G-H). On the test day, when the shock was omitted, we detected robust release of dopamine and a small increase in glutamate release in response to the tone onset. The unexpected absence of the shock did not result in altered release of either of these two neurotransmitters (figure 2E, G). To explore the specificity of dopamine release in response to aversive foot-shocks within the LSS, we performed fear conditioning while measuring dopamine release from the NAcC. We found that dopamine was released in the NAcC in response to both aversive foot-shocks and their predictive cues (supplementary figure S3A-B). These results indicate that the dopamine dynamics in the LSS are similar to those in the NAcC. To test if dopamine signaling via D1-receptors is necessary for the activity of the MC4R-LSS neurons, we compared shock-induced calcium transients with and without D1-antagonism. Blocking D1-receptors did not decrease the size of the shock-induced peak in neural activity, while it decreased the movement of the animal during the session, proving pharmacological effect (figure 2I-L). These results suggest that MC4R-LSS neurons are activated by aversive foot-shocks and that this increase is not driven by D1-receptor signaling on the neurons.

### In the LSS, MC4R-neural activity and glutamate release, but not dopamine release, are correlated to movement

During the fear conditioning test, we observed a striking correlation between the neural activity and the movement of the animal, measured in pixels changed per video frame, which includes all movements done by the animal (figure 3A). A cross-correlation of the signals revealed a high correlation with virtually no time-shift (figure 3B), indicating that MC4R-LSS neurons are a part of the movement selection and initiation system. To investigate whether the movement correlation is specific to MC4R-LSS neurons, we measured GcaMP6s signal from non-specific neurons in the lateral part of the nucleus accumbens core (NAcC) during the same conditions. We found that the neural activity measured from all neurons in the NAcC is also highly correlated to the movement of the animal (supplementary figure S3C), indicating that the correlation between neural activity and body movements is not specific to MC4R-neurons nor the LSS. In contrast to the neural activity of MC4R-LSS neurons, dopamine release measured with fiber photometry was not correlated to the movement of the animal when measured in either the LSS or the NAcC (figure 3C-D, supplementary figure S3D), suggesting that the neural activity of MC4R-LSS neurons is not directly driven by dopamine. Glutamate release in the LSS shared dynamics with MC4R-LSS-neural activity during fear conditioning and was also correlated to animal movement (figure 3E-F), suggesting that glutamate is the fast driver of MC4R-LSS neurons.

**Figure 3.**
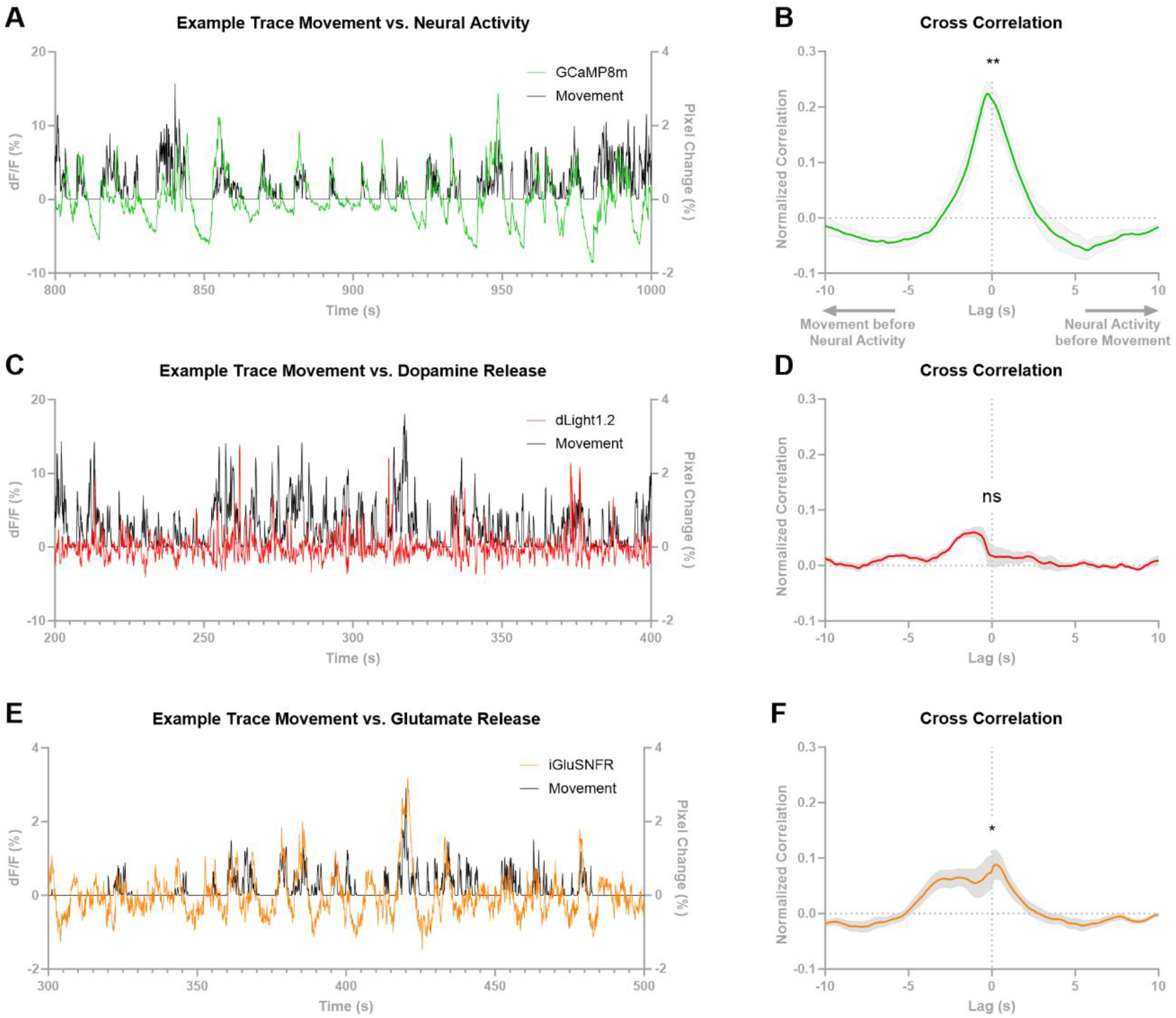
MC4R-LSS-neural activity is correlated to body movement. (A) Part of an example trace of MC4R-lateral stripe of the striatum (LSS) neuronal activity measured with fiber photometry and the calcium sensor GCaMP8m plotted together with body movement of the animal (measured as pixel change per video frame during the fear test) showing a correlation between the two measurements. (B) A cross-correlation of neural activity and movement shows a positive correlation with almost no time shift. R-values at lag 0 are significantly different from 0 (n=6, one sample t-test). (C) Part of an example trace of dopamine release in the LSS measured with fiber photometry and the dopamine sensor dLight1.2 plotted together with body movement of the animal showing that dopamine release in the LSS is not correlated to animal movement. (D) A cross-correlation of dopamine release and movement showing that dopamine release is not correlated to the movement of the animal (n=7, one sample t-test). (E) Part of an example trace of glutamate release onto D1-LSS-neurons measured with fiber photometry and the glutamate sensor iGluSNFR plotted together with body movement of the animal showing a correlation between glutamate release and movement of the animal. (F) A cross correlation of glutamate release and movement shows a correlation with no time shift 0 (n=10, one sample t-test). Collectively, these data indicates that MC4R-LSS neurons are mainly driven by glutamate, not dopamine, in the short term. *p < 0.05; **p < 0.01.

### MC4R-LSS neurons are activated by sweet rewards

Having established the responsiveness of MC4R-LSS neurons to aversive foot-shocks, we further investigated their response to rewarding stimuli and compared it with aversion-induced neural activation. In a Pavlovian conditioning paradigm, food-restricted mice with a genetically encoded calcium indicator (GcaMP8m) and an optic fiber placed above the LSS were trained using a tone predicting the delivery of a sugar pellet. MC4R-LSS neurons were activated by sweet rewards but not by the predictive tone (figure 4A-C, supplementary video V1). Unpredicted and predicted pellets generated equal responses, and the omission of a predicted pellet did not affect the neural activity in any direction (supplementary figure S4A-B). To compare the magnitude of neural responses induced by rewarding and aversive stimuli, we exposed the trained animals to a combined session consisting of 10 sugar pellet deliveries followed by 10 aversive foot-shocks. In line with previous fear conditioning experiments, the MC4R-LSS neural activity increased in response to foot shocks (figure 4C). Comparison of the peak post-stimulus calcium signal reveals that the activation of MC4R-LSS neurons induced by aversive and rewarding stimuli are of similar magnitude (figure 4D).

**Figure 4.**
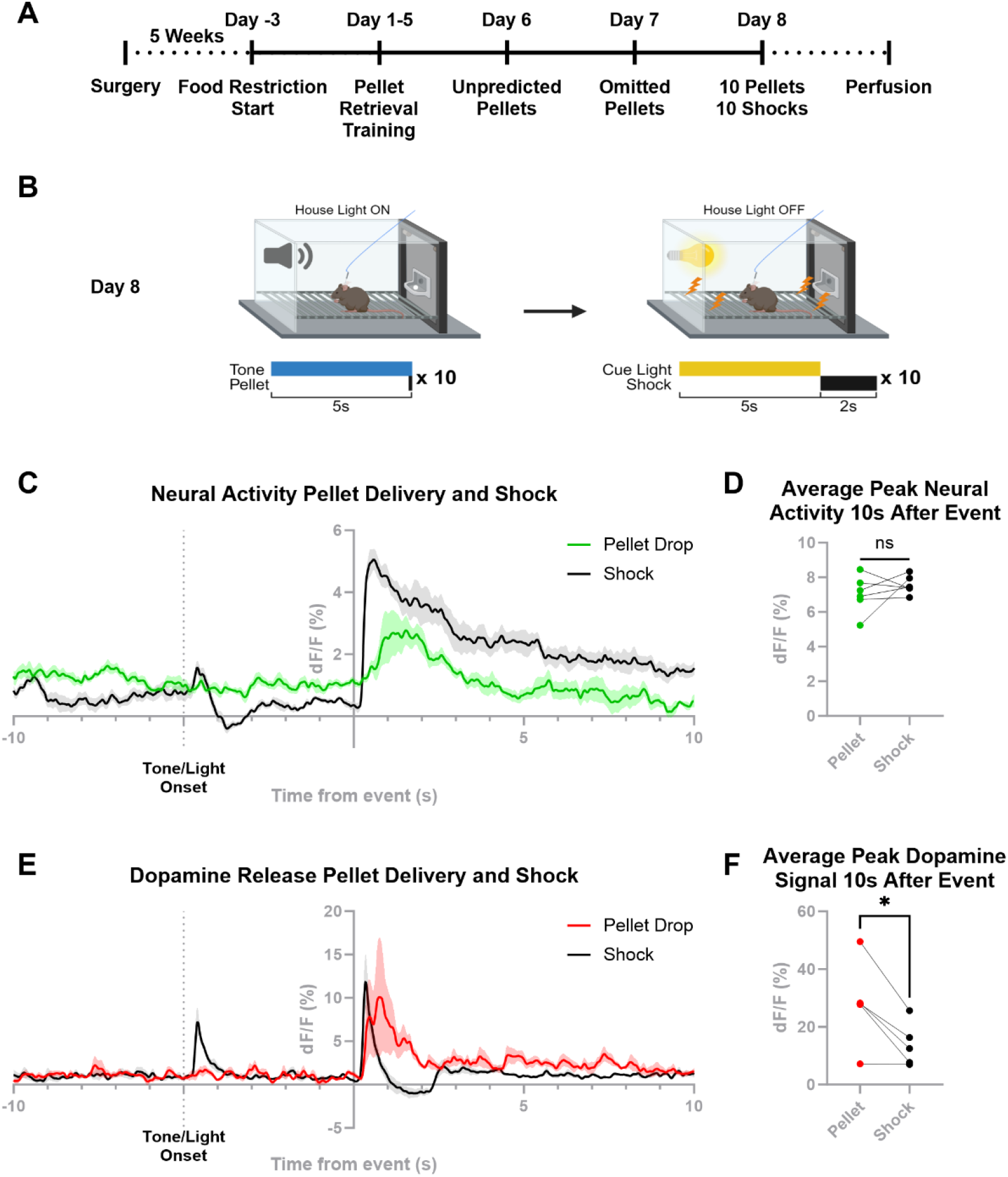
MC4R-LSS neurons are activated, and dopamine is released in the LSS in response to sugar pellet consumption. (A) Timeline of the experiment. (B) Trial structure of the combined reward and aversion session on day 8. (C) Mean data from the combined reward and aversion session on day 8. Neural activity is increased in response to both pellet consumption and to aversive footshocks (mean from 10 trials for each stimulus per animal, n=6 mice). (D) Maximal peak height within 10s after pellet delivery or footshock. Calcium signal magnitude is similar in response to pellet consumption and footshock (mean from 10 trials for each stimulus per animal; n=6 mice, two-tailed paired t-test). (E) Dopamine measurements (dLight1.2) from LSS during the same paradigm as in (C). Dopamine is released in response to both pellet consumption and foot shock. No response is seen to the tone cue predicting pellet delivery. Dopamine is released in response to the light cue predicting footshock (mean from 10 trials for each stimulus per animal, n=5 mice). (F) Maximal dopamine release 10s after the event is higher in response to pellet drop compared to foot-shock (mean from 10 trials for each stimulus per animal, n=5 mice, two-tailed paired t-test). *p < 0.05.

### Dopamine is released in the LSS during consumption of a sweet reward

Since dopamine is released in response to aversive stimuli, we tested the dopamine response in the LSS during the consumption of a sweet reward. Food-deprived mice were exposed to the same rewarding Pavlovian conditioning paradigm as described above. Dopamine release was measured in the LSS using a dLight 1.2 sensor and fiber photometry. Dopamine release was evident in the LSS when the mice consumed the pellet, but no response was observed to the onset of the predictive tone (figure 4E, supplementary video V2). We could not see any reward prediction error encoding by dopamine release in the LSS, since unpredicted and predicted pellets generated equal responses, and the omission of a predicted pellet did not affect the dopamine release in any direction (supplementary figure S4C-D). To compare the magnitudes of aversion- and reward-induced dopamine transients, trained mice were tested in a session comprising of 10 pellet deliveries followed by 10 foot-shocks paired with cue-lights. As seen in the previous fear-conditioning experiment, a robust dopamine release occurred in response to both foot-shock and cue onset (figure 4E). Further comparison revealed that the reward-induced dopamine release was larger than the aversion-induced release (figure 4F). Notably, during the behavioral sessions, we observed that dopamine was released in response to both the onset and offset of the house light in the operant chamber, suggesting that dopamine is released in the LSS in response to salient stimuli (supplementary figure 4E-F).

### Activation of MC4R-LSS neurons is rewarding and increases locomotion

Since the striatum is highly involved in affective behaviors as well as motor initiation and given that the MC4R-LSS neurons are activated by both rewarding and aversive stimuli, we next investigated the valence and locomotor effect of optogenetic activation of these neurons (Figure 5A-C). We found that in an intracranial self-stimulation task, hChR2-expressing mice, but not EYFP-expressing controls, were highly motivated to nose-poke for unilateral optogenetic activation of MC4R-LSS neurons (figure 5D). The same hChR2-expressing mice exhibited a preference for the light-paired chamber in a real-time preference paradigm, whereas the control group was indifferent to light stimulation (figure 5E). We next tested exploratory behavior in an open field box. Activation of MC4R-LSS neurons for 5 min caused a robust increase in locomotion (figure 5F). Next, we investigated whether the rewarding effect of activating MC4R-neurons was specific to the LSS or if similar activation in other parts of the striatum would generate similar results. Activation of MC4R-expressing neurons in the NAcC produced similar behavioral effects (supplementary figure S5A-C) but no rewarding effects were observed by activating MC4R-neurons in the medial nucleus accumbens shell (supplementary figure S5D-E) or the dorsal striatum (supplementary figure S5F-G). To assess whether MC4R-LSS neurons are necessary for voluntary locomotion, food intake, and energy homeostasis, we tested the impact of ablation of these neurons on locomotor behavior in an open field, voluntary wheel-running, food intake and body weight. Ablation of the MC4R-LSS neurons with Caspase 3 did not result in a decrease in locomotion or wheel-running behavior, nor did it affect food intake or body weight (supplementary figure S6). These results suggest that the MC4R-LSS neurons are part of the reward system and are involved in motor control, but that they are not necessary for voluntary movement, wheel-running behavior, or food intake.

**Figure 5.**
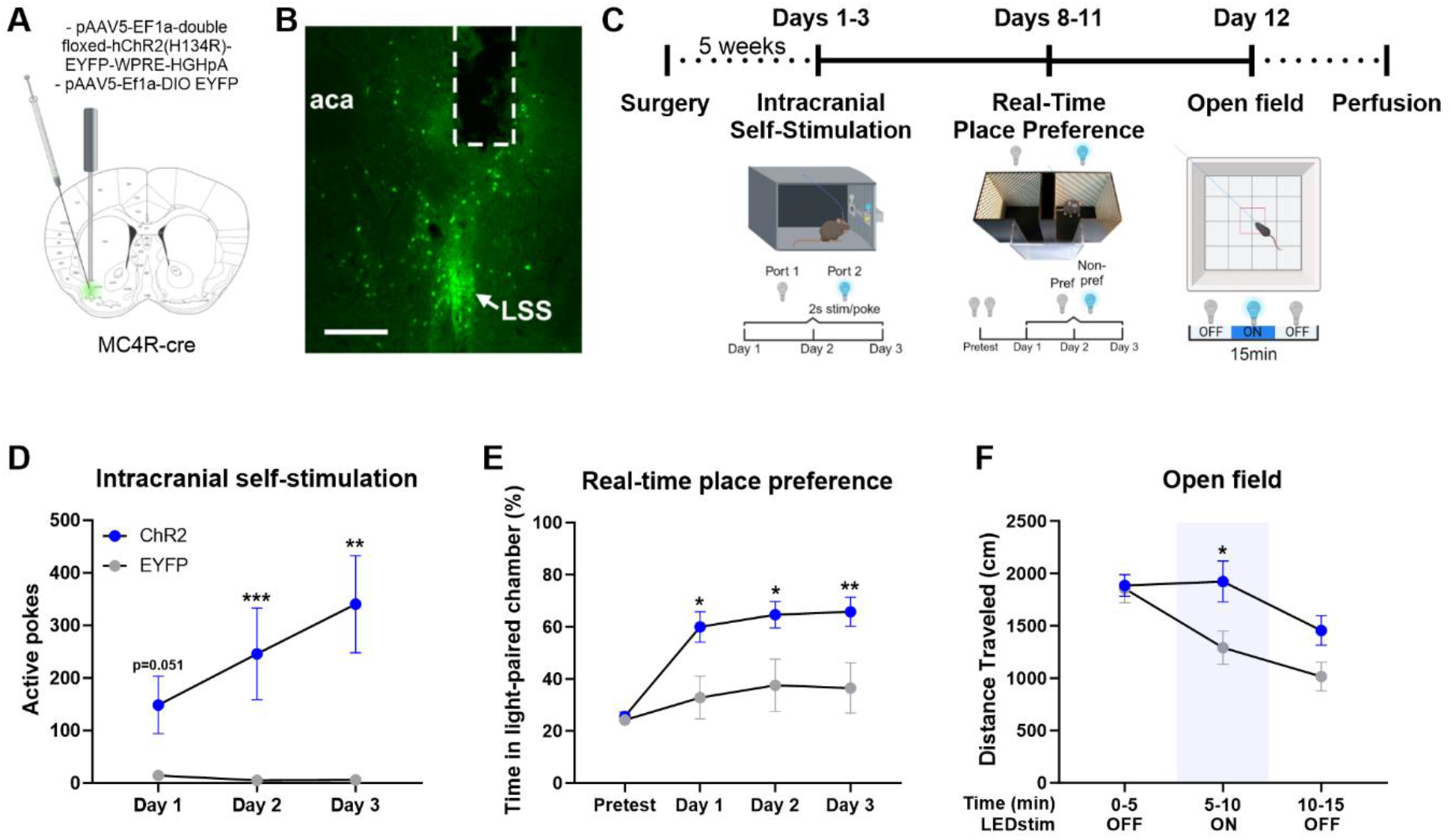
Optogenetic activation of MC4R-LSS neurons is rewarding and increases locomotion. (A) Experimental design for photoexcitation of MC4R-expressing neurons in the lateral stripe of the striatum (LSS). (B) Representative image of a coronal striatal slice expressing ChR2 in MC4R-LSS neurons. (C) Timeline of experiments. (D) Mice learn to self-stimulate for unilateral activation of MC4R-LSS neurons in an optogenetic intracranial self-stimulation paradigm (n=9 ChR2, n=7 EYFP, multiple two-tailed Mann-Whitney comparison tests corrected for multiple comparisons with False Discovery Rate with the method of Two-stage step-up (Benjamini, Krieger, and Yekutieli), ChR2 vs. EYFP). (E) The same mice with ChR2 expression in MC4R-LSS neurons prefer a chamber paired with light stimulation, contrary to DIO-EYFP controls (n=9 ChR2, n=7 EYFP, repeated-measures ANOVA followed by Šídák’s multiple comparisons test, ChR2 vs. EYFP). (F) Open field locomotion before, during and after 5 min pulsed unilateral optogenetic stimulation of MC4R-LSS neurons. Data is shown per 5-minute time bins (n=9 ChR2, n=7 EYFP, repeated-measures ANOVA followed by Šídák’s multiple comparisons test, ChR2 vs. EYFP). *p < 0.05; **p < 0.01; ***p < 0.001.

### Activation of MC4R-LSS neurons increases local dopamine release in the LSS

D1-neurons in the lateral nucleus accumbens have been shown to increase dopaminergic cell firing via disinhibition in the lateral VTA (Yang et al., 2018). Using a dual optogenetic approach, we co-injected the constitutive dLight1.2 sensor and the cre-dependent, red-shifted opsin rsChrmine (or an mScarlet control virus) into the LSS of MC4R-cre mice. This allowed us to simultaneously measure dopamine release and manipulate MC4R-LSS-neuronal activity with blue and red light, respectively, through a chronically implanted fiber. Optogenetic activation of MC4R-LSS neurons led to a robust and dose-dependent increase in dopamine release in the LSS. The magnitude and duration of this dopamine response scaled with the light intensity and stimulation duration, respectively. No effect on dopamine release was seen in the mScarlet control group except for a mild response to very high light powers in some mice (figure 6A-F). To test whether this dopamine release is mediated by a local circuit in the striatum, we conducted an ex-vivo dual optogenetic experiment using striatal coronal slices. First, we validated that MC4R-LSS neurons could be activated in the setup by co-injecting viral vectors encoding cre-dependent GcaMP8m and rsChrmine in the LSS of one hemisphere and GcaMP8m and an mScarlet as a control in the other hemisphere of MC4R-cre mice. Stimulation of rsChrmine with red light induced robust and repeatable increases in GcaMP8m signal while no increase in GcaMP8m signal could be measured from the mScarlet control slices, indicating that MC4R-LSS neurons can be activated by red-shifted optogenetics in a coronal slice (figure 6G). Further, coronal slices from MC4R-cre mice injected with constitutive dLight1.2 and cre-dependent rsChrmine in the LSS of one hemisphere and a dLight1.2 and an mScarlet control in the other were tested. While spontaneous dopamine releases could be measured in the slice, no dopamine release was observed in response to the activation of MC4R-LSS neurons, suggesting that the feedback loop is not mediated by a local striatal circuit (figure 6H, supplementary figure S7).

**Figure 6.**
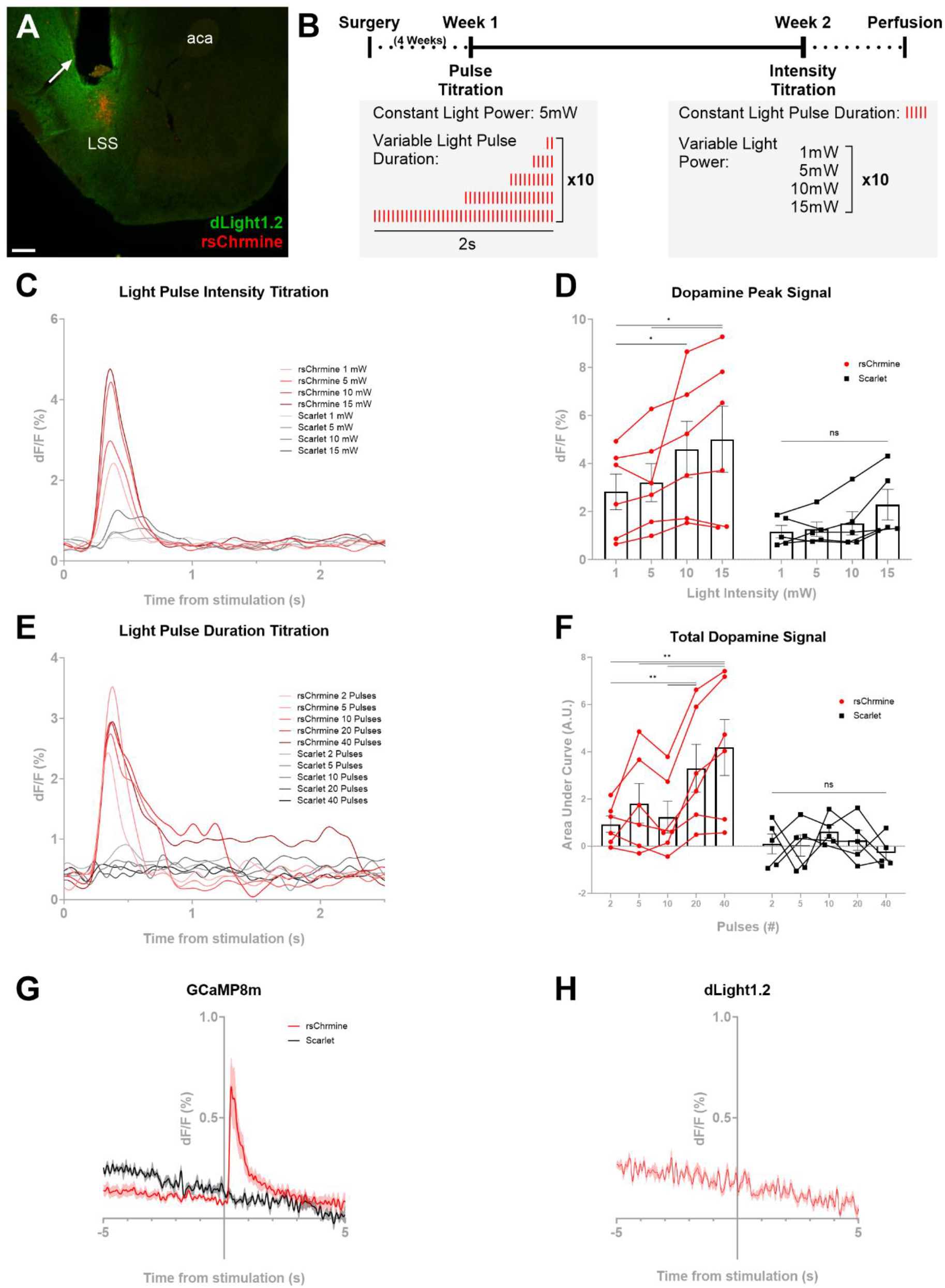
Activation of MC4R-LSS neurons causes a dose-dependent dopamine release in the lateral stripe of the striatum (LSS). (A) Representative image of a coronal striatal slice expressing constitutive dLight1.2 and rsChrmine-mScarlet in MC4R-LSS neurons. Fiber tract marked with arrow. Scale bar 200 µm. aca: anterior commissure. (B) Timeline of dual optogenetics and fiber photometry experiment and trials. (C) Traces from light intensity titration show a positive dose-response relationship between red light optogenetic activation of MC4R-LSS neurons expressing rsChrmine and dopamine release in the LSS. (D) Same data as in (C) displayed as maximal dF/F within 1 second following stimulation showing a significant dose-response relationship between peak dopamine release and light intensity in the rsChrmine group but not in the mScarlet controls (repeated-measures ANOVA followed by Šídák’s multiple comparisons test). (E) Increased stimulation duration of MC4R-LSS neurons increases the duration of elevated dopamine in the LSS. (F) Same data as in (E) expressed as area under the curve during 3s after stimulation start showing a significant dose-response relationship between total dopamine release and light pulse duration in the rsChrmine group but not in the mScarlet controls (n=6 rsChrmine, n=5 Scarlet controls; repeated-measures ANOVA followed by Šídák’s multiple comparisons test). (G) MC4R-LSS neurons in a coronal slice can be activated by the red-shifted opsin rsChrmine. Traces showing increased GCaMP8m signal from MC4R-neurons during optogenetic stimulation while the GCaMP8m signal from control slices is unaffected (n=16 rsChrmine slices from 4 animals, n=6 control slices from 2 animals). (H) Same as in (G) but for dLight1.2 measurements. No dopamine release was seen in response to optogenetic stimulation of MC4R-LSS neurons, indicating that there is no local feedback-loop (n=7 slices from 3 animals). *p < 0.05; **p < 0.01.

## Discussion

Here, we present a functional characterization of MC4R-expressing neurons in the LSS. Our findings demonstrate that these neurons are activated by both rewarding and aversive stimuli, while the activation of this population is rewarding. In addition, we measured dopamine and glutamate release in the LSS during the same behavioral paradigms to compare neural activity with afferent neurotransmitter release, indicating that glutamate is the fast driver of MC4R-LSS neuronal activity. Our results also show a positive feedback loop from striatal medium spiny neurons to dopamine release within the same area.

Lately, multiple single-cell RNA sequencing papers have highlighted the heterogenicity among striatal medium spiny neurons and identified atypical D1-neurons. One subpopulation of atypical D1-neurons is enriched in the LSS and expresses high levels of Tshz1 and FOXP2 (Chen et al., 2021; Gokce et al., 2016; Saunders et al., 2018; Stanley et al., 2020). We found that the LSS is enriched in MC4R-expressing neurons, and that they express D1-receptor and FOXP2, indicating that most MC4R-LSS neurons are atypical D1-neurons. Consequently, the MC4R-cre mouse line is an attractive tool to selectively target atypical D1-neurons of the LSS. Whereas the MC4-LSS neurons were atypical at the transcriptional level, our optogenetics findings indicate that their behavioral function is similar to that of typical D1 neurons in that activation is rewarding and induces locomotion (Cole et al., 2018; Kravitz et al., 2010; Soares-Cunha et al., 2016). This contrasts the previous finding that activation of Tshz1-expressing cells (another marker of atypical D1-neurons) in the striosomes in the dorsal striatum was aversive (Xiao et al., 2020). The fact that activation of MC4R-neurons in the NAcC was also rewarding could indicate that the same neuronal type found in the LSS is present in the entire ventrolateral striatum, or that there are two or more distinct populations of MC4R-expressing neurons with ability to induce reward.

We found that MC4R-LSS neurons were activated by both aversive and rewarding stimuli, suggesting that they detect salient events independent of valence. Although the MC4R-LSS neurons were activated also by aversive stimuli, their activation was strongly rewarding. This might suggest that these neurons could, under certain circumstances, assign positive value to salient stimuli. Alternatively, there might be several sub-populations of MC4R-LSS neurons activated by different stimuli to induce distinct affects. The counterintuitive feature of being activated by aversive signals while mediating reward might relate to the finding that mice lacking MC4Rs develop place preference to both aversive and rewarding stimuli, a phenotype that could be rescued by MC4R re-expression in D1 neurons in the striatum (Klawonn et al., 2018). It is possible that the MC4R-neurons in the striatum are overactive, or in another way given excessive weight in the valence assigning process in mice lacking MC4Rs. This would go in line with a study showing long-term depression of striatal D1 neurons after alpha-MSH stimulation (Lim et al., 2012). However, it is unclear if the lack of MC4Rs affects the excitability of MC4R-LSS neurons. It is also unclear if, and in such case how, the MC4Rs in the LSS are activated in the context of aversion. MC4R-expression is found in many brain areas spared from hypothalamic POMC-neurons afferents (Lima et al., 2019), and here we show that this is also the case for the LSS. It is possible that MC4R in the LSS can be activated by volume transmission of alpha-MSH or circulating Lipocalin 2 – which has been shown to be an MC4R-agonist (Mosialou et al., 2017). Since the MC4R has a high constitutive activity (Kleinau et al., 2020), it is possible that inverse agonists like agouti-related peptide could affect neural activity without the presence of an agonist. Alternatively, other, yet unidentified ligands could be the main effector of MC4R in the LSS.

In this study, we describe neural activity of MC4R-LSS neurons and dopamine release in the LSS during two different Pavlovian conditioning paradigms, one aversive and one rewarding. While foot-shocks and pellet consumption triggered both neural activity peaks and dopamine release, there were several discrepancies between the two measurements. Dopamine was released in response to cues predicting aversive stimuli, while the MC4R-neurons did not show overt activity in response to conditioned stimuli. In the rewarding Pavlovian conditioning paradigm, we did not observe any dopamine response to the conditioned tone, despite extensive training, while dopamine response to cues predicting aversive outcomes was developed rapidly. This might be explained by the more salient and time-locked features of the foot-shock in the fear conditioning compared to the pellet drop, where the animal can collect the pellet at any time after delivery. Alternatively, dopamine release in the LSS is more responsive to aversion prediction than to reward prediction. This would be surprising, however, since the dopamine release in response to reward consumption was higher than in response to the aversive foot-shock.

Dopamine release in the striatum has been described as a saliency signal (Kutlu et al., 2021, 2023; Yawata et al., 2023). In line with this finding, we found that stimuli with seemingly limited value, such as the house light of the operant chamber turning on or off caused robust dopamine release in the LSS. However, although the initial dopamine response to an aversive foot-shock was positive, the dopamine signal was lower than baseline at the end of the shock. This could indicate a mixed response from dopaminergic terminals, where one population increases the firing due to salience and the other is inhibited by negative value. The dopamine release in response to foot-shock was not LSS-specific, since the same pattern could be seen in the NAcC. These findings contrast previous studies suggesting that aversive stimuli lead to a decrease in striatal dopamine in all subregions (van Elzelingen et al., 2022). Possibly, this discrepancy could be explained using different dopamine measurement techniques (fast-scan cyclic voltammetry versus photometry) and different aversive stimuli, where the foot-shock we delivered is likely to be more salient than the white noise used in the aforementioned study. Since the by-far largest dopamine responses recorded in our study were in response to consumption of rewarding sugar pellets, dopamine could have a dual role of saliency signaling and reward signaling in the LSS and NAcC, possibly via heterogeneous dopaminergic neural populations (De Jong et al., 2022).

Dopamine and the striatum are well known for their involvement in motor functions. Here, we found a clear correlation between body movements of the animal, including those not leading to locomotion, and the activity of MC4R-LSS neurons. Moreover, artificial activation of these neurons increased locomotion. This strongly suggests that the neurons are involved in movement control and highlights the importance to consider the impact of all body movements when studying the striatum. The fact that ablation of the MC4R-LSS neurons did not affect locomotion could be explained by the small numbers of neurons affected, compared to the whole striatum, as the movement correlation is not LSS- or MC4R-specific. Whereas MC4R-LSS-neural activity was correlated to movement in the short timescale, we show that the dopamine release in the LSS did not. Even though dopamine has been shown to drive neural activity with sub-second resolution *ex vivo* (Lahiri & Bevan, 2020), this was not the case in our *in vivo* experiments. Rather, the discrepancies in neural activity and dopamine release during Pavlovian conditioning experiments and the observation that the administration of a D1-antagonist did not attenuate the foot-shock-induced neural activity strongly implies that the fast, neural activity driving input to striatal neurons is not dopamine. Likely, the main driver of acute changes in MC4R-LSS-neuronal activity is glutamatergic cortico-striatal inputs, with dopamine as a slow modulator, since glutamate is released in the LSS in a pattern similar to the activity of MC4R-LSS neurons. While dopamine is not the primary driver of activity of the MC4R-LSS neurons, we instead show a causal relationship between activity of MC4R-LSS neurons to dopamine release in the LSS. This is likely mediated via disinhibition in the midbrain, since there are strong projections from MC4R-LSS neurons to the substantia nigra pars reticulata. Moreover, no dopamine release was seen when MC4R-LSS neurons were activated in a coronal slice, where these projections were severed. Our finding that stimulating MC4R-LSS neurons leads to dopamine release in the stimulated area *in vivo* complements a previous electrophysiological study showing that D1-neurons in the ventrolateral striatum can increase the activity of midbrain dopaminergic neurons through disinhibition (Yang et al., 2018).

In conclusion, MC4R-LSS neurons are primarily atypical D1-neurons that are activated by both aversive and rewarding stimuli, while their activation is rewarding and induces locomotion. The feature of detecting salience, regardless of the valence, and inducing positive affect might contribute in the positive direction when assigning value to salient stimuli under some conditions.

### Limitations of the study

Even though we used a MC4R-cre transgenic mouse line and stereotactic surgeries to specifically target MC4R-LSS neurons, both optogenetic and fiber photometry techniques have inherent limitations in spatial specificity. While the placement of the optical fibers ensured that the majority of the targeted neurons were located in the LSS, the size of the fibers relative to the small size of the LSS precludes us from excluding the possibility that MC4R-neurons adjacent to the LSS contributed to the behavioral effects or neural responses we observed using these techniques. Our optical methods also lack cellular resolution. Thus, it could be that we missed more subtle divisions of MC4R-LSS neurons into subclusters with different functional roles. A second limitation of our study is the apparent lack of behavioral effects of chronic ablation of MC4R-LSS neurons in the caspase experiment. Indeed, this manipulation neither affected locomotion in the open field test or the wheel running experiment, nor did it alter food intake. On the other hand, we observed clear hyperlocomotion upon optogenetic activation of this neuronal cluster. One interpretation of these contrasting findings could be that although the activity MC4R-LSS neurons can modulate motor behavior, it is not necessary for such behavior. Alternatively, compensatory mechanisms following the chronic ablation of these neurons may have occluded any behavioral effects. Future studies could use acute inhibition of these neurons with opto- or chemogenetic techniques and/or other behavioral readouts to better understand the physiological functions of MC4R-LSS neurons. Finally, it should be noted that although we established a role for MC4R-LSS neurons in affect regulation and motor control, the functional role of MC4R signaling in these processes was not addressed in this study.

## Acknowledgements

We thank the staff at the animal facility for technical assistance regarding animal breeding and care. We thank Maria Ntzouni and Vesa Loitto from histology and imaging facilities at Linköping Microscopy Unit for technical assistance. We also thank Larri Mohell Malinen, Johannes Sjölin and Sertan Arkan for their technical assistance, and Ilona Szczot for her help with the development of the fiber photometry analysis software. Finally, we thank the Technical Platform for Optogenetics and Optical Imaging from the Center for Systems Neurobiology at Linköping University, as well as Markus Heilig, for their role in establishing the local infrastructure for optogenetics and fiber photometry experiments. Part of the figures were created with biorender.com.

## Author contributions

Conceptualization, DE, GPL, JS and JW; Methodology, DE, GPL, JS and JW; Software Programming, JW; Validation, GPL and JS; Formal Analysis, GPL and JS; Investigation, GPL, JS and MN; Resources, DE; Data Curation, DE, GPL, JS and JW; Writing – Original Draft Preparation, GPL and JS; Writing – Review & Editing Preparation, DE, GPL, JS and JW; Visualization Preparation, GPL and JS; Supervision, DE and JW; Project Administration, DE; Funding Acquisition, DE and GPL.

