## Supplemental figures for "Melanocortin 4 receptor-expressing neurons in the lateral stripe of the striatum are involved in affect regulation and motor control"

**Supplementary Figures**

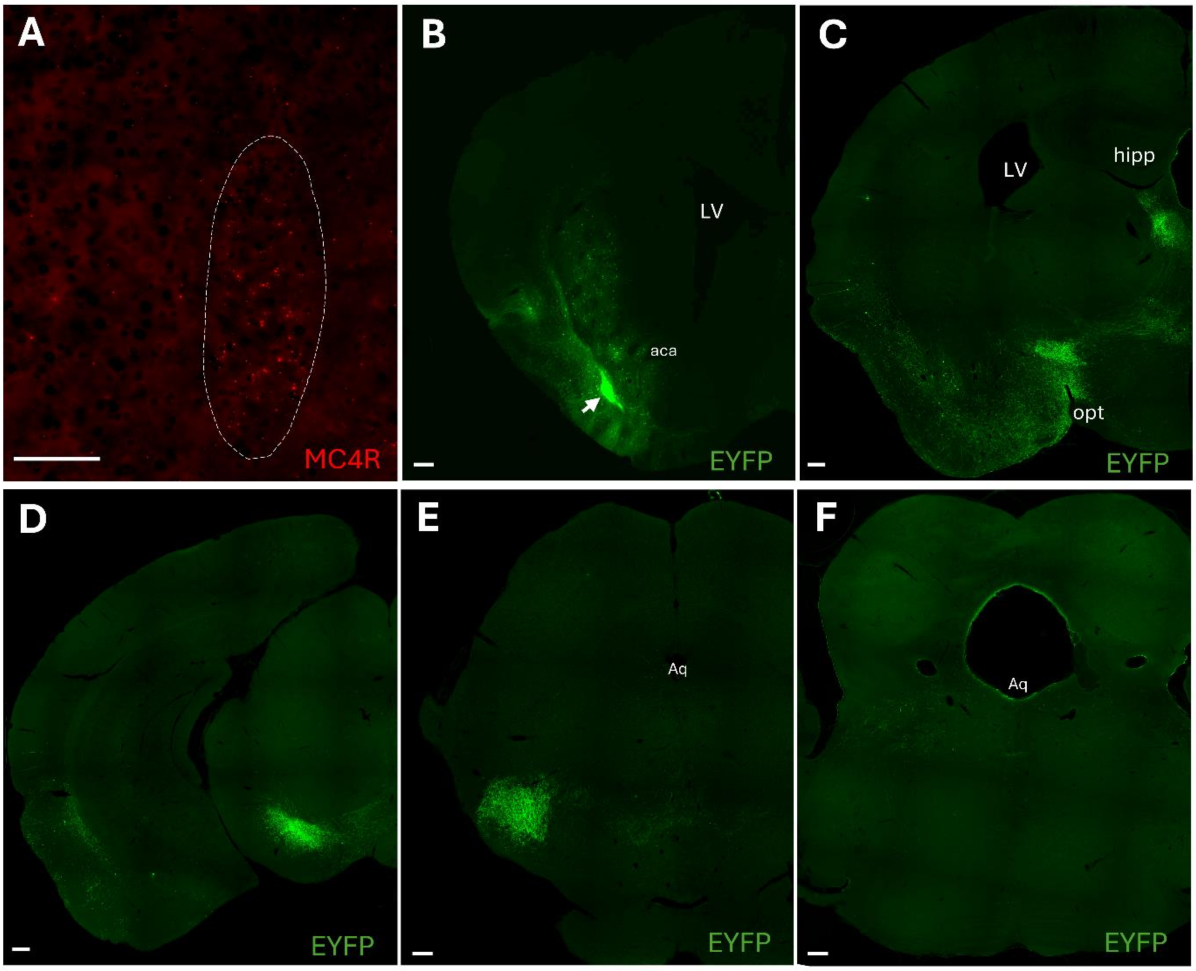


*Supplementary figure S1. MC4R-neurons are present in the adult lateral stripe of the striatum and project to the substantia nigra pars reticulata.*

(A) Fluorescent in situ hybridization of a coronal section from an adult mouse showing ongoing MC4R-expression in the lateral stripe of the striatum (LSS). LSS marked with dashed line. Scale bar 100 µm. (B-F) Projection mapping from MC4R-LSS neurons. Injection of 100 nl AAV loaded with cre-dependent EYFP in the LSS. Selection of rostral-caudal sections stained with immunohistochemistry against EYFP. Expression of EYFP is seen at the injection site (B; arrow) and in projections to the lateral hypothalamus, epithalamus (C) and midbrain. Strong immunostaining can be seen in the substantia nigra pars reticulata (D-F). Scale bars 200µm. LV: Lateral ventricle, aca: anterior commissure, hipp: Hippocampus, opt: optic tract, aq: Aqueduct.

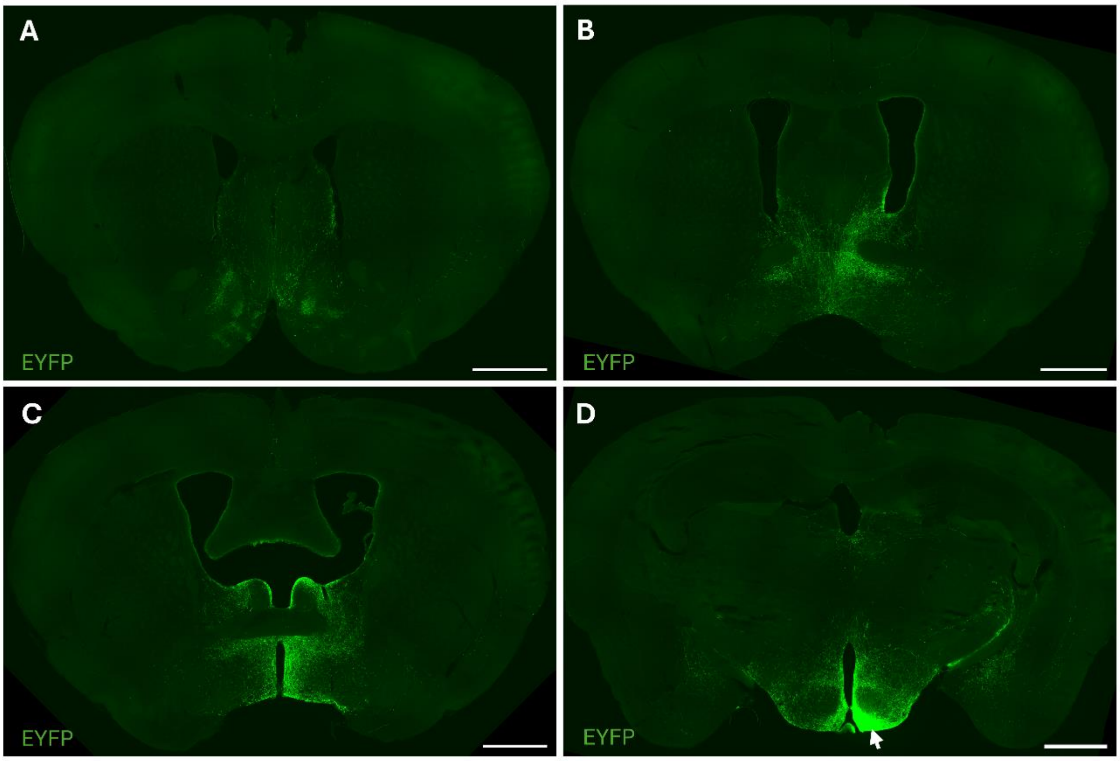


*Supplementary figure S2. No projections from arcuate nucleus POMC-neurons were found in the lateral stripe of the striatum.*

(A-D) Rostral-caudal images from a POMC-cre mouse injected with cre-dependent EYFP in the arcuate nucleus. Projections are seen in the medial nucleus accumbens shell and other areas, but not in the lateral stripe of the striatum (LSS) or the dorsal striatum. Injection site shown in D (arrow). Scale bars 200μm.

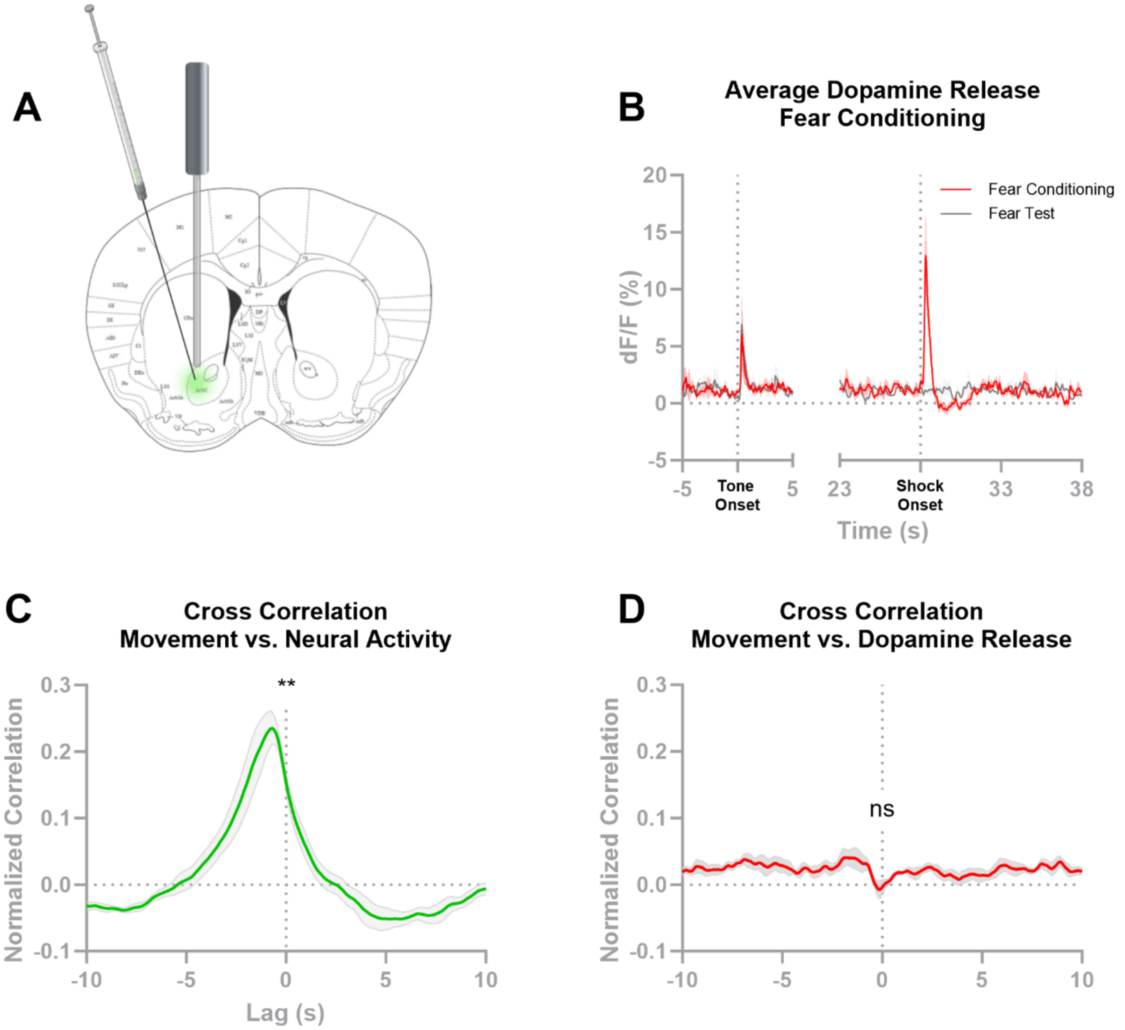


*Supplementary figure S3. Dopamine is released in the nucleus accumbens core by aversive footshocks and their paired cues.*

(A) Schematic showing site of injection and fiber placement in the nucleus accumbens core (NAcC). (B) Dopamine release measured with dLight1.2 from the NAcC during fear conditioning and fear test the following day. Dopamine is released in response to aversive footshocks and their predictive tone (n=4). (C) Cross-correlation of neural activity of unspecified neurons of the NAcC, measured with constitutive GCaMP6s, and body movements of the mouse during fear test. A clear correlation is seen between neural activity and body movements. R-values at lag 0 are significantly different from 0 (n=3, one sample t-test). (D) Cross-correlation of dopamine release in the NAcC and body movements of the mice during fear test show that dopamine release is uncorrelated to movement of the animal R-values at lag 0 are not significantly different from 0 (n=4, one sample t-test). **$p < 0.01$

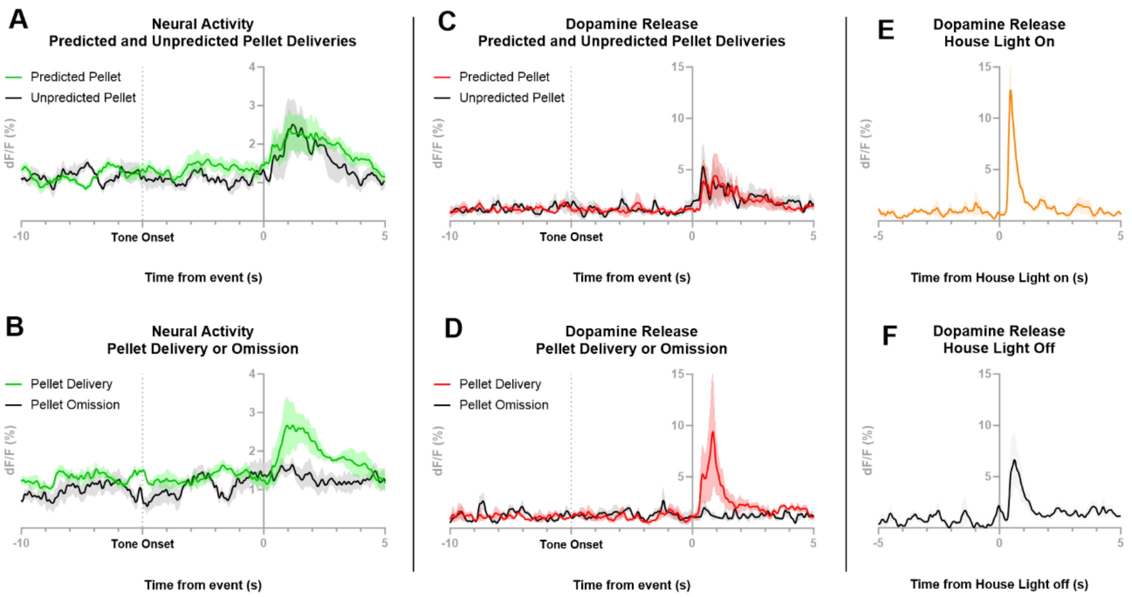


*Supplementary figure S4. No reward prediction error encoding by MC4R-expressing neurons or dopamine in the lateral stripe of the striatum.*

(A) The neural response, measured with GCaMP8m, is similar after predicted and unpredicted pellets, suggesting there is no positive reward prediction error encoded by the MC4R-lateral stripe of the striatum (LSS) neurons. (B) The neural activity is not decreased by omission of the predicted reward, suggesting there is no negative reward prediction error encoded by MC4R-LSS neurons (n=6). (C-D) Same as in (A-B), but for dLight1.2 measurements of dopamine release. (C) Similar dopamine release was seen in response to predicted and unpredicted pellet deliveries, suggesting there is no positive reward prediction error encoded by dopamine release in the LSS. (D) The dopamine release is not decreased by omission of the predicted reward, suggesting there is no negative reward prediction error encoded by dopamine release in the LSS (n=5). (E) Dopamine is released in the LSS in response to house light turning on (mean from 5 training sessions, n=5). (F) Dopamine is released in the LSS in response to house light turning off (Mean from 5 training sessions, n=5).

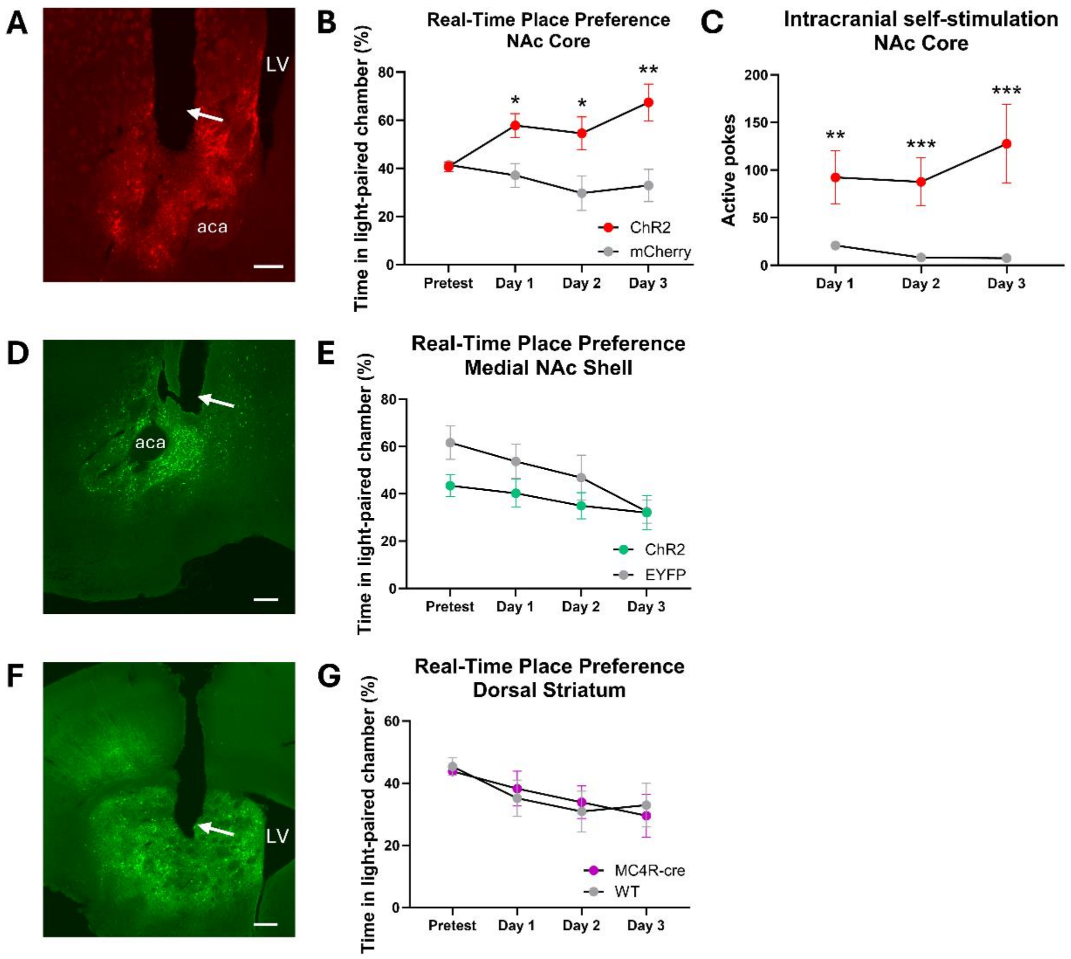


*Supplementary figure S5. Activating MC4R-neurons in the nucleus accumbens core, but not in the medial nucleus accumbens Shell or in the dorsal striatum, is rewarding.*

(A) Representative picture showing viral expression (pAAV5-EF1a-double floxed-hChR2(H134R)-mCherry-WPRE-HGHpA) and fiber placement of MC4R-cre animals injected in nucleus accumbens (NAc) core. The arrow marks the fiber tract. (B) Mice with opsin expression (ChR2) in nucleus accumbens core neurons prefer a chamber paired with light stimulation, contrary to EYFP controls (n=12 ChR2, n=10 mCherry, repeated-measures ANOVA followed by Šídák's multiple comparisons test, ChR2 vs. mCherry). (C) Mice self-stimulate to optogenetically activate MC4R-neurons in the nucleus accumbens core (n=16 ChR2, n= 12 mCherry controls, multiple two-tail Mann-Whitney comparison test, ChR2 vs. mCherry). (D) Representative picture showing viral expression (pAAV5-EF1a-doublefloxed-hChR2(H134R)-EYFP-WPRE-HGHpA) and fiber placement of MC4R-cre animals injected in medial nucleus accumbens shell. The arrow marks the fiber tract. (E) No effect was seen in a real-time place preference paradigm in response to optogenetic activation of MC4R-neurons in the medial nucleus accumbens shell (n=8 ChR2, 6 EYFP controls, repeated-measures ANOVA followed by Šídák's multiple comparisons test, ChR2 vs. EYFP). (F) Representative picture showing viral expression (pAAV5-EF1a-doublefloxed-hChR2(H134R)-EYFP-WPRE-HGHpA) and fiber placement of MC4R-cre animals injected in Dorsal Striatum. The arrow marks the fiber tract. (G) No effect was seen in a real-time place preference paradigm in response to optogenetic activation of MC4R-neurons in the Dorsal Striatum (n=8 MC4R-cre, 7 wildtype (WT) littermate controls, repeated-measures ANOVA followed by Šídák's multiple comparisons test, MC4R-cre vs. WT). Scale bars 200 µm. LV: Lateral ventricle, aca: anterior commissure. *$p < 0.05$; **$p < 0.01$; *** $p < 0.001$.

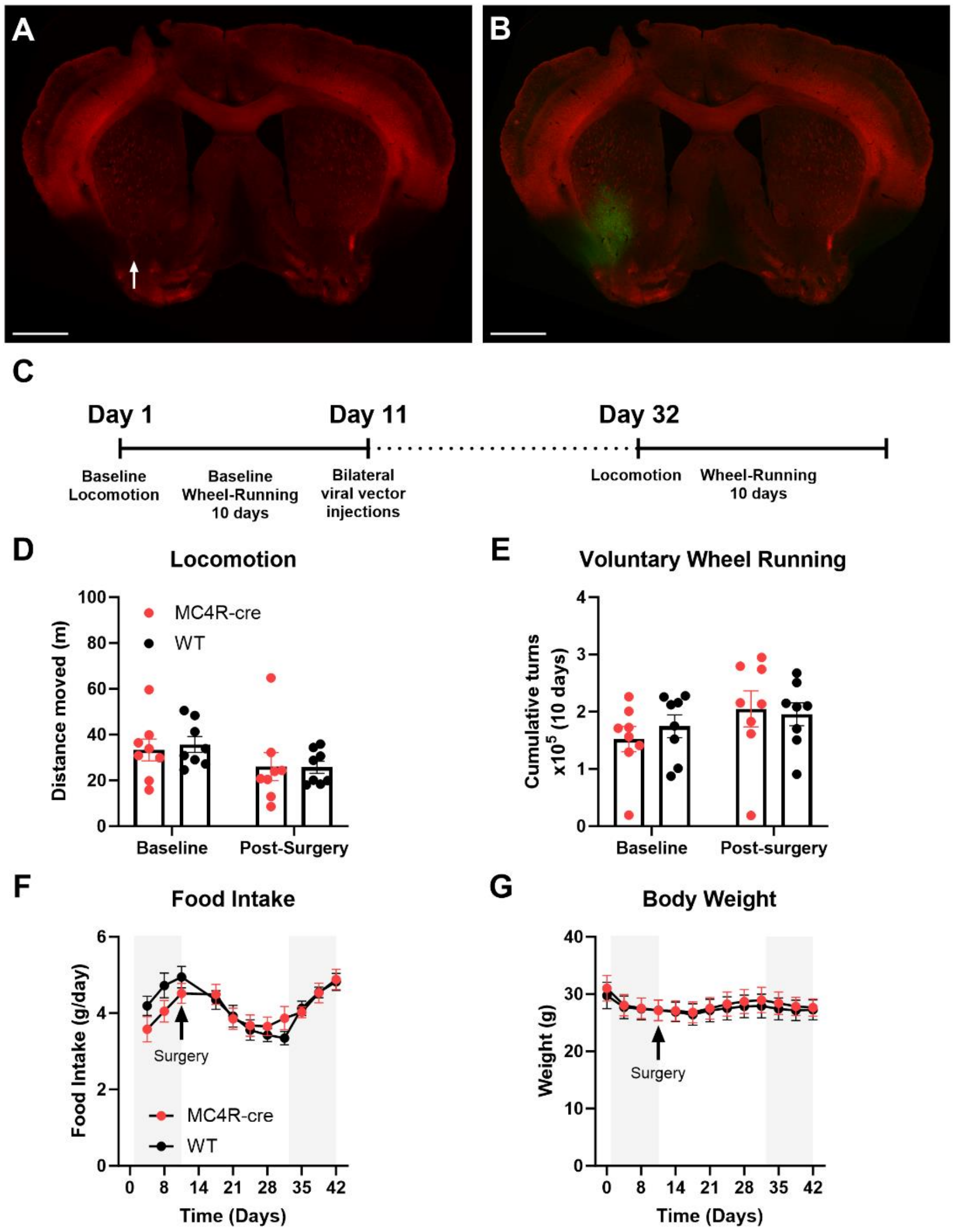


*Supplementary figure S6. Ablation of MC4R-LSS neurons does not affect locomotor or appetitive behaviors.*

(A-B) Validation of caspase3-ablation of MC4R-LSS-neurons. Representative picture of MC4R-cre-tdTomato mouse injected with cre-dependent caspase3 and constitutive EYFP in the lateral stripe of the striatum (LSS). Scale bars 1 mm. (A) No tdTomato-expressing MC4R neurons can be seen in the injected side (arrow) while the LSS is clearly visible on the control side. (B) Overlap with EYFP expression showing the spread of viral vector injection. (C) Timeline of experiment shown in (D-E). (D) Open field locomotion is not affected by ablation of MC4R-LSS neurons. (E) Ablation of MC4R-LSS neurons does not affect voluntary wheel-running behavior. (F) Food intake was measured every 3-4 days during the experiment. Data expressed as grams/day. No difference was seen between mice with ablated MC4R-LSS neurons compared to littermate controls. (G) No difference in body weight seen before or after ablation of MC4R-LSS neurons compared to wildtype littermates. Shaded areas in (F-G) indicate time in wheel-running boxes. n=8 per group.

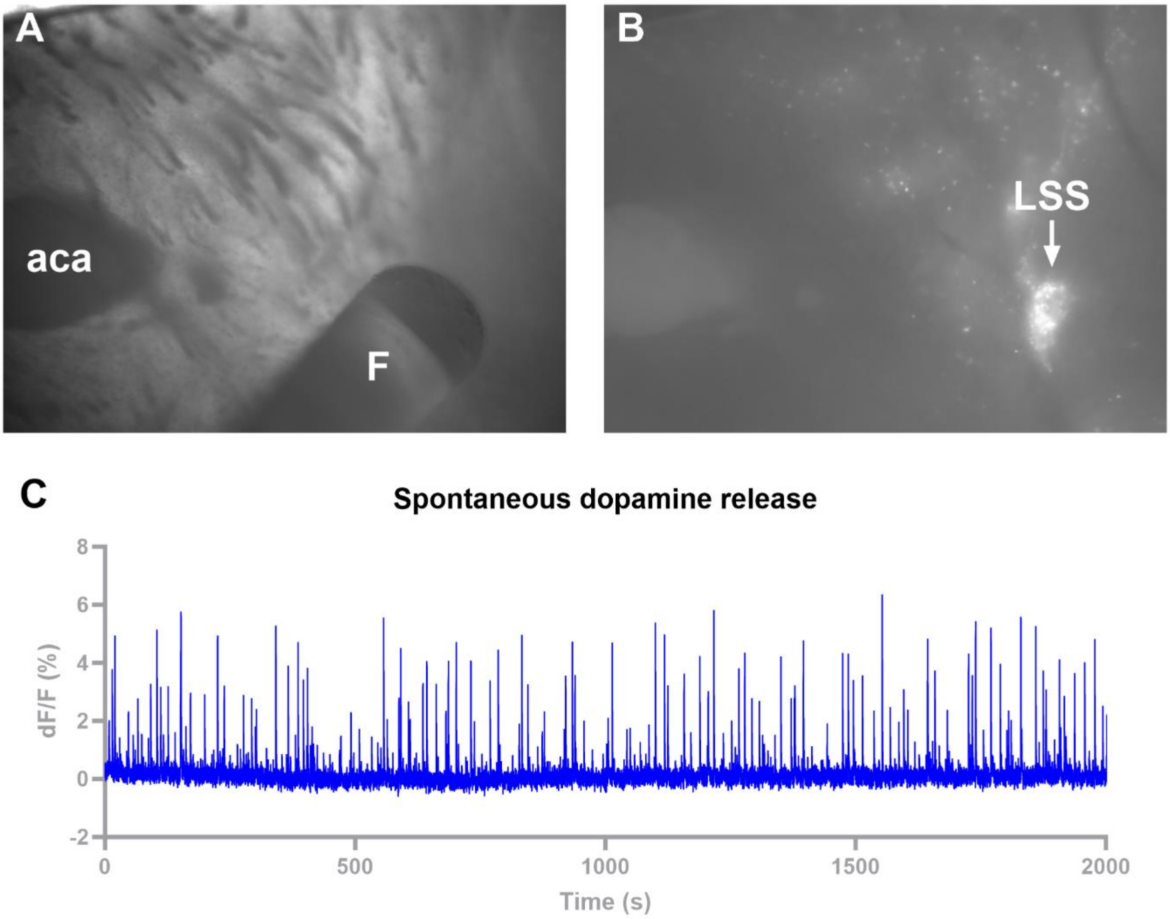


*Supplementary figure S7. Spontaneous dopamine release can be measured from a coronal slice using fiber photometry.*

(A) Picture showing fiber placement over the lateral stripe of the striatum (LSS) in a coronal slice of the striatum. aca: a commissure, F: optic fiber. (B) Expression of rsChrmine in the LSS in the same coronal slice as in (A). (C) Example graph showing spontaneous dopamine peaks in a coronal slice proving that dopamine release occurs and can be measured under the experimental conditions.

*Supplementary video V1. MC4R-LSS neurons are activated by consumption of a rewarding sugar pellet.*

Example video showing GCaMP8m measurements from MC4R-neurons in the lateral stripe of the striatum (LSS) aligned with video from day 8 of experiment shown in figure 4b. MC4R-LSS neurons are activated by consumption of a sugar pellet.

*Supplementary video V2. Dopamine is released in the LSS by consumption of a rewarding sugar pellet.*

Example video showing dLight1.2 measurements from the lateral stripe of the striatum (LSS) aligned with video from day 8 of experiment shown in figure 4D. Dopamine is released in the LSS by consumption of a sugar pellet.
